# The modular origin and evolutionary expansion of the enigmatic DGF-1 protein family in trypanosomatids

**DOI:** 10.64898/2026.07.27.741050

**Authors:** Mathias J. Mangino, Juan Manuel Trinidad-Barnech, Adriana Parodi-Talice, Fernando Alvarez-Valin, Luisa Berná

## Abstract

Large multigene families are a hallmark of trypanosomatid genomes and play major roles in genome plasticity and parasite adaptation. Among them, the Dispersed Gene Family 1 (DGF-1) of *Trypanosoma cruzi* remains one of the least understood because of its large size, repetitiveness, and fragmented annotations. Here, using a high-quality long-read assembly of *T. cruzi*, we performed comprehensive re-annotation and structural characterization of DGF-1. We identified seven tandemly repeated structural modules, here termed *ribs*, that form the extracellular region. This is followed by a distinct C-terminal membrane-associated region containing a conserved *hat* domain and multiple transmembrane helices. These domains are highly conserved across paralogous copies and the 7 *ribs* cluster according to positional identity rather than gene origin, indicating that the canonical seven-rib architecture predates the expansion of the family. DGF-1 proteins are present in the early-branching trypanosomatid species *Paratrypanosoma confusum* and several other trypanosomatids but absent in *Leishmania* and African trypanosomes, consistent with multiple independent secondary losses. Comparative analyses across Euglenozoa indicate that the DGF-1 architecture did not originate in free-living bodonids. Instead, *Bodo saltans* contains proteins with rib-like domains and others containing the membrane-associated region, whereas the first complete DGF-1 architecture appears in *P. confusum*. Phylogenetic and comparative genomic analyses indicate that the family subsequently underwent lineage-specific losses and independent expansions, the largest expansion occurring in *T. cruzi*. Together, these findings reconstruct the evolutionary emergence of the DGF-1 architecture from pre-existing structural modules and provide a framework for one of the largest and most enigmatic gene families in trypanosomatids.

## Introduction

Large lineage-specific expansions of gene families are a hallmark of trypanosomatid genomes and are thought to underlie the remarkable adaptability of these parasites to diverse hosts and environments. In *Trypanosoma cruzi*, the causative agent of Chagas disease, this genomic expansion is particularly pronounced, with approximately half of the genome composed of repetitive sequences, including large multigene families and retrotransposons (Balouz et al. 2025; Berná et al. 2018; El-Sayed et al. 2005; Reis-Cunha et al. 2022). Many of these families encode surface or secreted proteins involved in host–parasite interactions, immune evasion, and cell invasion, and are frequently located in nonsyntenic regions of the genome, which are associated with increased genomic plasticity (Buscaglia et al. 2006; De Pablos y Osuna 2012).

Among these expanded repertoires found in the *T. cruzi* genome, the Dispersed Gene Family 1 (DGF-1) represents one of the least understood examples (Ramírez 2023). DGF-1 genes encode unusually large proteins and are present in hundreds of copies across the genome. Early estimates based on the CL Brener reference *T. cruzi* strain suggested the presence of several hundred DGF-1 genes and numerous pseudogenes (El-Sayed et al. 2005; Wincker et al. 1990). More recent genome assemblies have revealed substantial variation in copy number across strains (Balouz et al. 2025; Berná et al. 2018; Greif et al. 2026; Hoyos Sanchez et al. 2024; Wang et al. 2021). Members of this family are distributed throughout the genome and have been detected in higher concentration in regions proximal to telomeres (Kim et al. 2005; Balouz et al. 2025; Cruz-Saavedra et al. 2025; Ramirez 2020), although a comprehensive characterization of their genomic distribution is still lacking. Phylogenetic and comparative analyses indicate that DGF-1 evolution is shaped by gene duplication, recombination, and gene conversion, while maintaining an overall high degree of sequence conservation with localized regions of variability (Kawashita et al. 2009a). In addition, the large number of pseudogenes may contribute to sequence diversification by serving as a reservoir of genetic variation.

Despite its abundance, peculiar structure (one of the largest proteins) and its atypical evolutionary dynamism, the biological role of DGF-1 remains poorly understood. The presence of structural features such as epidermal growth factor–like domains and RGD (Arg-Gly-Asp) motifs have led to the hypothesis that these proteins may participate in host–parasite interactions (De Pablos y Osuna 2012). However, experimental evidence indicates that at least some DGF-1 proteins display stage-specific expression, with higher abundance in amastigotes, challenging their classification as canonical surface proteins (Lander et al. 2010). These observations suggest that DGF-1 proteins may fulfill other roles beyond classical host–parasite adhesion.

From an evolutionary perspective, DGF-1 differs from other gene families expanded in *T. cruzi*, such as trans-sialidases, GP63 and mucins, which are typically organized into quite divergent subgroups (Dean et al. 2025; Berná et al. 2025; Freitas et al. 2011). In contrast, DGF-1 copies display a high degree of sequence conservation across paralogs, suggesting a more constrained and coordinated evolutionary regime (Kawashita et al. 2009b). Previous studies have identified localized regions of variability within DGF-1 proteins, but its overall amino acid sequence conservation is a signature of purifying selection, indicating that this family does not follow the same diversification patterns observed in most families of surface proteins (Kawashita et al. 2009a; Ramírez 2023).

However, the evolutionary relationships within DGF-1 remain incompletely resolved, in part due to limitations in genome assembly quality and gene annotation, particularly in repeat-rich genomic regions. The exceptionally large size of DGF-1 genes, together with their high sequence similarity across multiple genomic copies, complicates accurate genome assembly and gene annotation, potentially obscuring patterns of conservation and modular organization.

Here, we use a high-quality long-read assembly of the *T. cruzi* Dm28c genome to reconstruct the evolutionary history of the DGF-1 family. We integrate manual gene curation, structural characterization, domain-based homology searches, and comparative analyses across Euglenozoa to curate the complete DGF-1 repertoire, characterize its modular architecture, and trace the evolutionary history of its constituent domains. By integrating sequence- and structure-based approaches, we show how the DGF-1 protein architecture emerged very early and subsequently diversified across trypanosomatids, providing a new evolutionary framework for understanding one of the largest and most enigmatic gene families in the *T. cruzi* genome.

## Results – Outline

### 1. Curation and re-annotation of the DGF-1 gene family in *T. cruzi* Dm28c

The DGF-1 family constitutes one of the largest multigene families in the *Trypanosoma cruzi* genome. However, its repetitive nature and the presence of numerous fragmented copies have recurrently complicated its annotation. To establish a reliable reference set, we used the manually curated annotation of 75 DGF-1 genes from the *T. cruzi* Dm28c genome generated by Berná et al. 2018. In addition, to analyze the genomic organization of DGF-1 loci along chromosomes, we performed a systematic re-annotation of the DGF-1 repertoire in the recently assembled *T. cruzi* Dm28c genome generated using PacBio HiFi sequencing, which resolves the genome into 32 chromosomes with separated haplotypes (Greif et al. 2026).

Using this assembly as a reference, DGF-1 loci were identified by homology searches and manually curated to distinguish intact genes from fragmented copies and pseudogenes. We identified 64 putatively intact DGF-1 genes in haplotype H1 and a comparable number in H2 (62), together with 74 DGF-1 pseudogenes (Supplementary Table S1).

DGF-1 genes are notably long, with intact copies reaching a mean length of 10,242 bp. In contrast, pseudogene lengths display a bimodal distribution, with one group of short pseudogenes (fragments) and another approaching the length of intact genes (Supplementary Figure S1). However, many pseudogenes retain substantial sequence length and differ from intact genes primarily by disruptions in their coding sequence, including frameshifts caused by insertions or deletions and premature stop codons. Consequently, most pseudogenes were identified through nucleotide-based rather than protein-based homology searches (Supplementary Figure S1A).

We then examined the genomic distribution of DGF-1 loci along chromosomes in haplotype H1. Most chromosomes contain at least one DGF-1 locus, with the exception of three chromosomes in which no family members were detected (Figure 1A). Across the genome, the DGF-1 repertoire is relatively evenly distributed, with most chromosomes harboring between one and five loci. A notable exception is chromosome 12, which contains a large tandem cluster comprising 14 intact genes and three pseudogenes.

**Figure 1.**
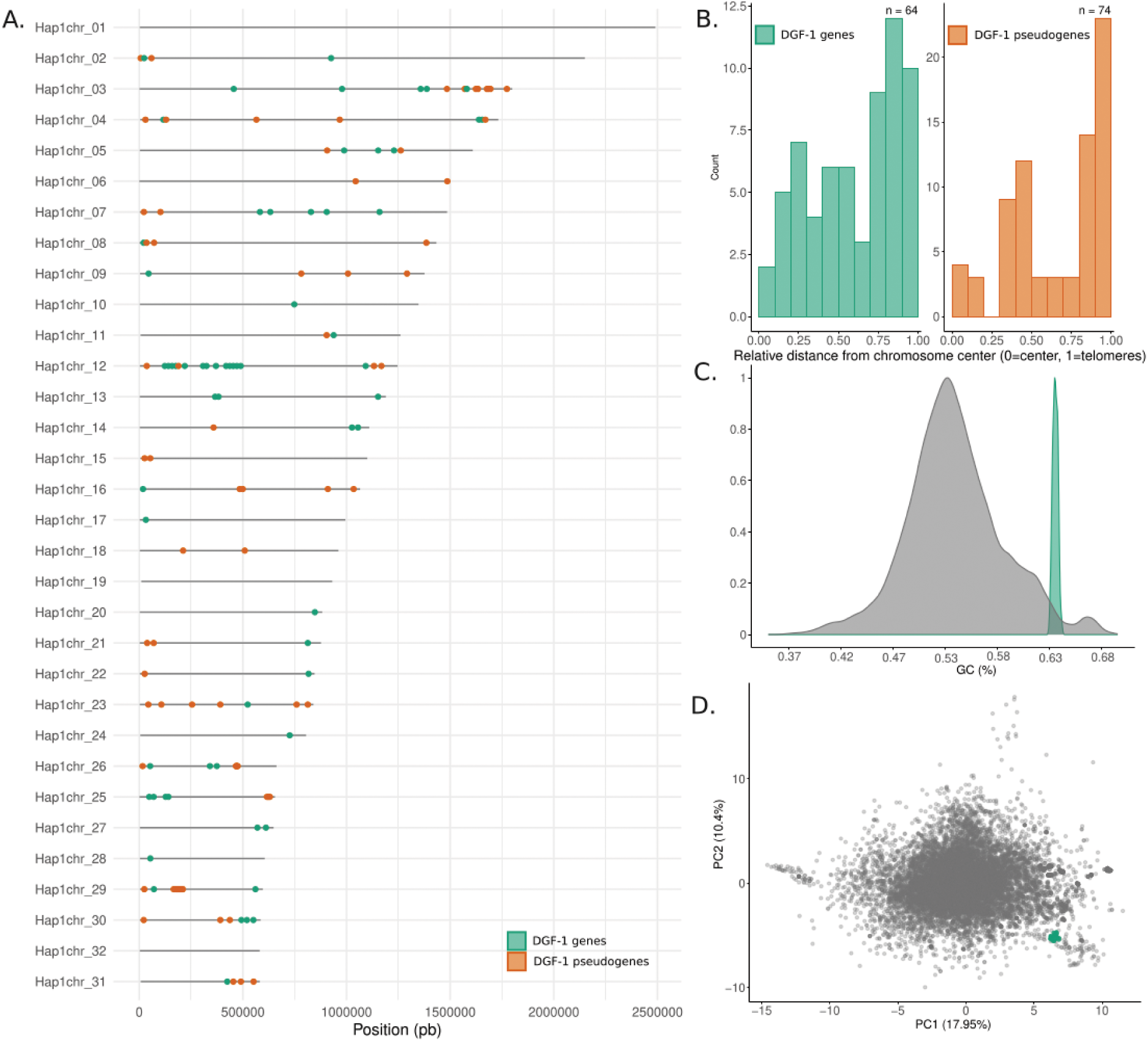
Genomic distribution and sequence composition of the DGF-1 family in *Trypanosoma cruzi* Dm28c. (A) Chromosomal distribution of the DGF-1 family across the 32 *T. cruzi* Dm28c chromosomes. Intact genes and pseudogenes are shown in green and orange, respectively. (B) Aggregated relative spatial distribution of DGF-1 genes (green) and pseudogenes (orange) across all chromosomes. Chromosome coordinates were normalized by distance to the nearest telomere, effectively collapsing both chromosomal ends into a single peripheral coordinate to represent gene density from the chromosome periphery toward the center. (C) Distribution of GC content across all coding sequences in the *T. cruzi* genome (gray) compared with that of the DGF-1 family (green). (D) Principal component analysis (PCA) of trinucleotide frequencies in coding sequences from the *T. cruzi* Dm28c genome, comparing all coding sequences (gray) with DGF-1 genes (green).

Both intact genes and pseudogenes are distributed along chromosomal arms but exhibit a clear bias toward chromosome ends, with an increased density in telomeric regions (Figure 1B). This pattern is evident from the relative positions of loci along normalized chromosome coordinates, where values approaching 1 correspond to telomeric regions. This not random spatial distributions of both intact genes and pseudogenes, differed significantly from those expected under a uniform distribution along the chromosome (Wilcoxon rank-sum test, *P* = 0.0045 and *P* = 3.1 × 10⁻⁵, respectively).

Besides their genomic distribution, DGF-1 genes exhibit a distinctive sequence composition that is highly homogenous among family members and differs markedly from the rest of the *T. cruzi* gene repertoire. This is reflected in their GC content, which shows a narrow distribution with a higher mean (63.8%) compared to the genome-wide average (51.3%) (Figure 1C, Supplementary Table S1). Consistently, principal component analysis based on trinucleotide composition reveals a clear segregation of DGF-1 genes, which cluster tightly in a distinct region of the compositional space, separate from the bulk of *T. cruzi* genes (Figure 1D). Together, these results indicate that DGF-1 genes are defined not only by their genomic organization but also by a distinctive compositional signature.

### 2. Phylogenetic relationships among DGF-1 copies reveal a gene expansion in *T. cruzi*

To assess sequence conservation within the DGF-1 family, we generated multiple sequence alignments of all curated H1 copies at both the nucleotide and amino acid levels and calculated pairwise sequence identities. DGF-1 genes are highly conserved, exhibiting a mean nucleotide identity of 88.3% (mean distance = 11.7%) and a mean amino acid identity of 84.2% (mean distance = 15.8%) across all pairwise comparisons. This overall conservation is readily apparent in the all-against-all amino acid identity matrix, which shows uniformly high similarity among most family members (Figure 2A).

**Figure 2.**
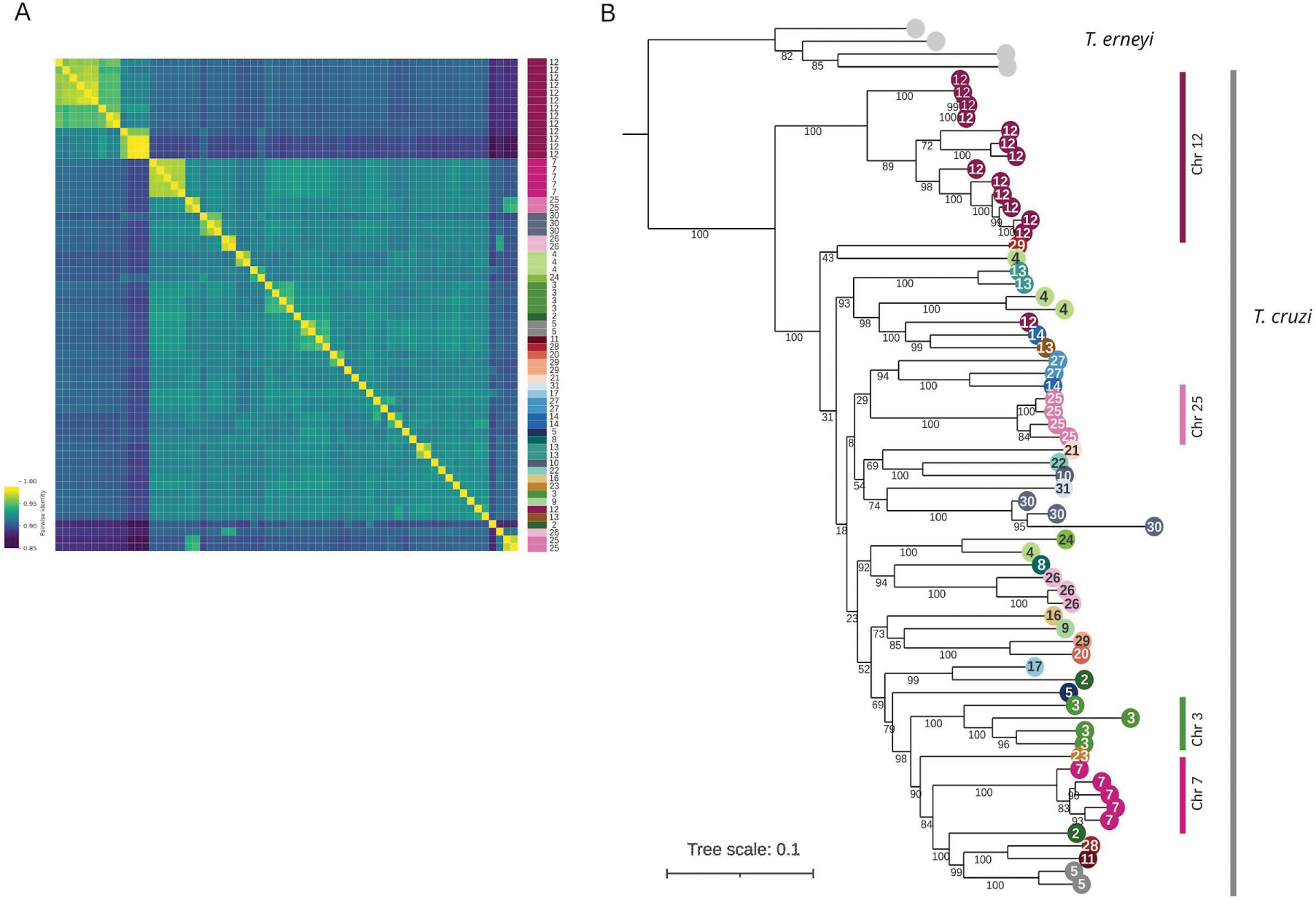
Phylogenetic relationships and pairwise sequence identity of the DGF-1 family. (A) Heatmap of pairwise amino acid sequence identities among DGF-1 variants, calculated from the multiple sequence alignment. Values range from dark blue (low identity) to bright yellow (high identity). (B) Rooted maximum-likelihood phylogenetic tree of DGF-1 amino acid sequences from *Trypanosoma cruzi* Dm28c strain. The tree was rooted using DGF-1 sequences from *Trypanosoma erneyi* as the outgroup, indicated by gray circles at the branch tips. *T. cruzi* sequences are color-coded according to their chromosome of origin.

Despite this high overall conservation, the identity matrix reveals several groups of closely related sequences. The most prominent cluster corresponds to the tandem array of 14 DGF-1 genes located on chromosome 12, while additional smaller clusters comprise genes from chromosomes 7 and 25. These relationships are further supported by the phylogenetic reconstruction, in which genes belonging to the same chromosomal clusters form.well-supported monophyletic groups (Figure 2B). Together, these results indicate that the DGF-1 family is highly homogeneous at the sequence level but contains subsets of paralogs that have undergone more recent local expansion and diversification.

To provide an evolutionary context for these relationships, we searched trypanosomatid genomes for homologous DGF-1 sequences and identified homologs in *Trypanosoma erneyi*. This trypanosome species is the closest relative of *T. cruzi, identified so far* (Lima et al. 2012; Cottontail et al. 2014). It is clearly located outside of immediate relatives like *T. cruzi marinkellei,* but much nearer than other species accepted to be close relatives to *T. cruzi*, like *T. lewisi* or *T. rangeli* (Bradwell et al. 2018). The DGF-1 protein encoding genes found in *T. erneyi* are very similar in length (3,450 amino acids) and have around 76% of amino acid identity with *T. cruzi* DGF-1 proteins. Rooting the phylogeny with this outgroup confirms that all *T. cruzi* DGF-1 copies form a single monophyletic group, indicating that the extensive expansion of the family occurred after the divergence of the *T. cruzi* and *T. erneyi* species.

To evaluate how the relationships among DGF-1 copies are presented outside the Dm28c reference genome, we incorporated DGF-1 sequences from additional *T. cruzi* strains belonging to distant lineages (Supplementary Figure S2). The resulting phylogeny yields the same major clades identified in Dm28c, indicating that the principal DGF-1 lineages emergence predate the diversification of extant *T. cruzi* strains. Rather than representing recent strain-specific duplications, these paralog groups have been maintained throughout the evolutionary history of the species.

Together, these findings indicate that the DGF-1 repertoire was established early during *T. cruzi* evolution and has remained remarkably stable despite the extensive duplication that characterizes the family. This evolutionary stability, combined with the high sequence conservation observed among intact copies, suggests that the major DGF-1 paralog lineages have been maintained during *T. cruzi* evolution.

### 3. Modular architecture of DGF-1 proteins in *T. cruzi*

To characterize the domain architecture of the DGF-1 family, we combined sequence-based annotation with protein structure prediction. Consistent with the high sequence similarity observed across DGF-1 copies (see above), InterProScan produced nearly identical domain profiles for all proteins (Figure 3A, Supplementary Figure S3 and Table S2). Given this conservation and the absence of structural models for full-length DGF-1 proteins, we selected a representative sequence for structure prediction using AlphaFold3. The resulting model was consistent with the InterProScan annotations while substantially improving domain resolution, allowing the identification of structural modules that could not be precisely delimited using sequence- or profile-based methods alone.

**Figure 3.**
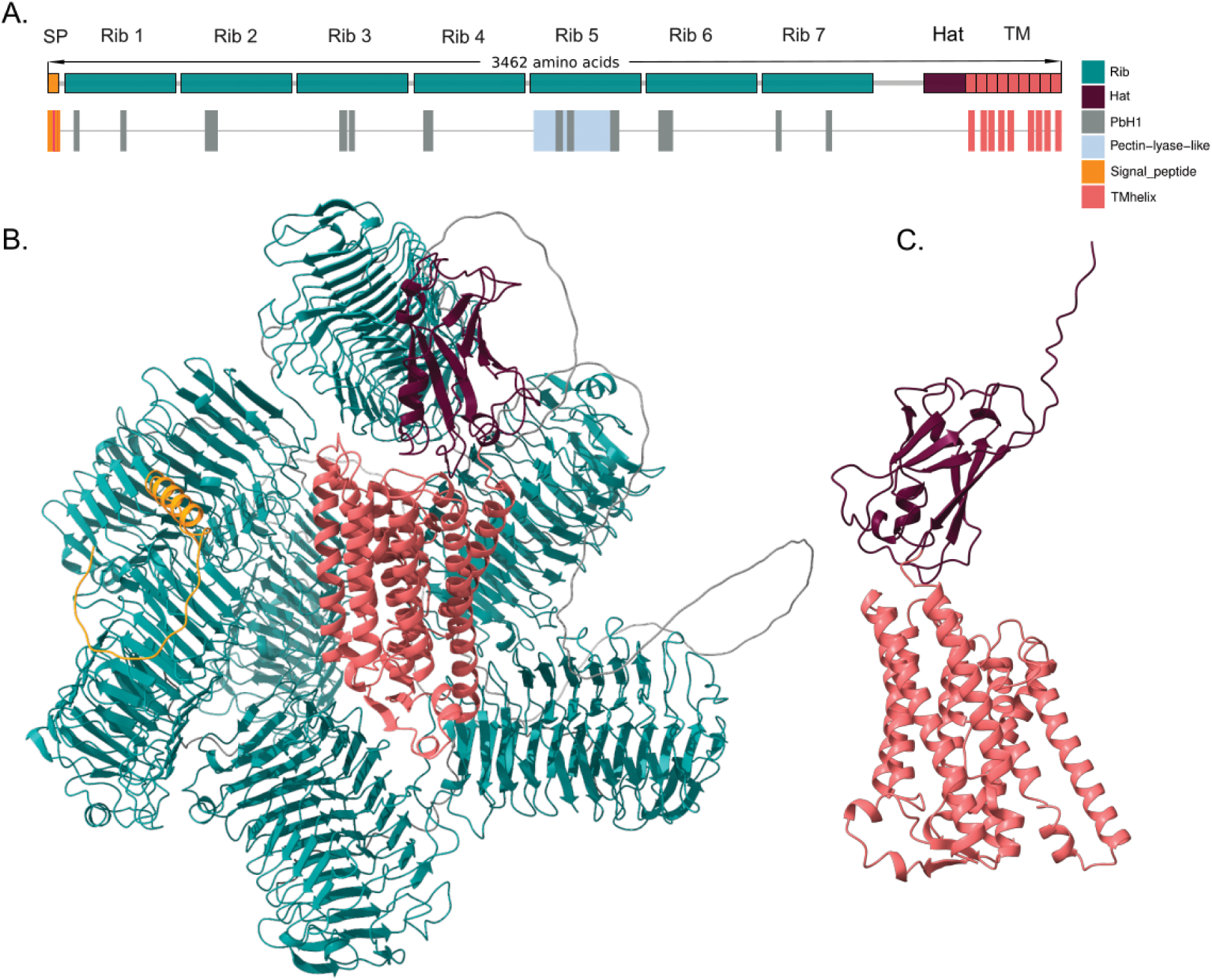
Structural and domain characterization of *T. cruzi* DGF-1 proteins. (A) Schematic representation of DGF-1 domain architecture. The top panel shows a linear representation based on the predicted 3D structure and topology, highlighting the signal peptide (orange), *rib* modules (teal), and the C-terminal region, which comprises the *hat* domain (wine) and the transmembrane helices (salmon). The bottom panel shows InterProScan domain predictions, including the signal peptide (orange), predicted transmembrane helices (salmon), PbH1 (gray; parallel beta-helix repeats, SM00710), and pectin lyase-like domains (light blue, SSF51126). (B) Predicted three-dimensional structure of a representative DGF-1 protein generated with AlphaFold3, shown as a ribbon model. Domains are colored according to panel A. (C) Close-up view of the C-terminal region.

The predicted structure reveals a highly conserved modular organization composed of a N-terminal signal peptide, followed by seven tandemly arranged structural modules, hereafter referred to as *ribs* modules, and a conserved C-terminal region consisting of a previously undescribed globular domain, designated the *hat* domain, followed by seven to nine transmembrane α-helix (Figure 3B and Supplementary Figure S4). Whereas the signal peptide and transmembrane helices are consistently detected by InterProScan, the remaining architecture only becomes fully apparent after structural prediction.

The *rib* modules constitute the most peculiar structural feature of DGF-1 proteins. Each module spans over approximately 390 amino acids and adopts a right-handed parallel β-helix fold, identified here for the first time in DGF-1 proteins. Each *rib* forms an elongated dextrorotatory solenoid composed of approximately twelve consecutive coils (Figure 3C and Supplementary Figure S5). AlphaFold3 predictions showed consistently high confidence for the individual *rib* modules, whereas the linker regions connecting consecutive *ribs* exhibited substantially lower confidence scores. Consequently, the relative arrangement of the *ribs* and the overall topology of the full-length protein remain uncertain and should be interpreted with caution.

Another domain identified as PbH1 in InterPro is found in all of the *ribs*, whereas a Pectin Lyase-Like profile consistently maps to specific portions of *rib* 5, and in some DGF-1 copies also in *rib* 3 (Supplementary Figure S3). Likewise, structural searches against the CATH database classified all *rib* modules within the single-stranded right-handed β-helix superfamily (CATH 2.160.20.10), consistent with the AlphaFold3 structural predictions. Some additional InterPro annotations were also detected within these regions, including EGF-like motifs. These annotations likely reflect similarities in local structural architecture rather than conservation of the biological functions typically associated with those protein families.

Immediately downstream of the seventh *rib*, we identified a compact domain of approximately 150 amino acids that was not detected by any sequence-based annotation method (Figure 3C, wine-colored domain). Structural similarity searches using Foldseek suggested that this previously unrecognized region adopts an antiparallel immunoglobulin-like β-sandwich fold. The highest-confidence structural matches were consistently assigned to CATH homologues superfamily 2.60.40.10, although the overall structural similarity remained moderate (Supplementary Figure S6). Given its characteristic position between the repetitive *rib* region and the transmembrane helices, we designate this previously undescribed module as the *hat* domain.

### 4. Structural and evolutionary relationships among *rib* modules

The remarkable structural similarity and conserved tandem organization of the *rib* modules suggest a shared evolutionary origin. To investigate their evolutionary relationships, we extracted all *rib* sequences from the 64 curated DGF-1 genes and reconstructed their phylogenetic history.

Phylogenetic analysis reveals that the 448 *rib* modules cluster according to their positional identity (Figure 4A). *Rib* modules also show high amino acid similarity within each positional class (mean pairwise identity among the same type of *rib* ∼83%; whereas *ribs* of different types have on average ∼30%, supplementary Figure S7), consistent with the clustering observed in the phylogenetic analysis. Specifically, *rib* 1 sequences from different DGF-1 copies group together, as do *rib* 2, *rib* 3, and so on. This pattern indicates that each *rib* position represents a distinct module that is conserved across paralogous genes.

**Figure 4.**
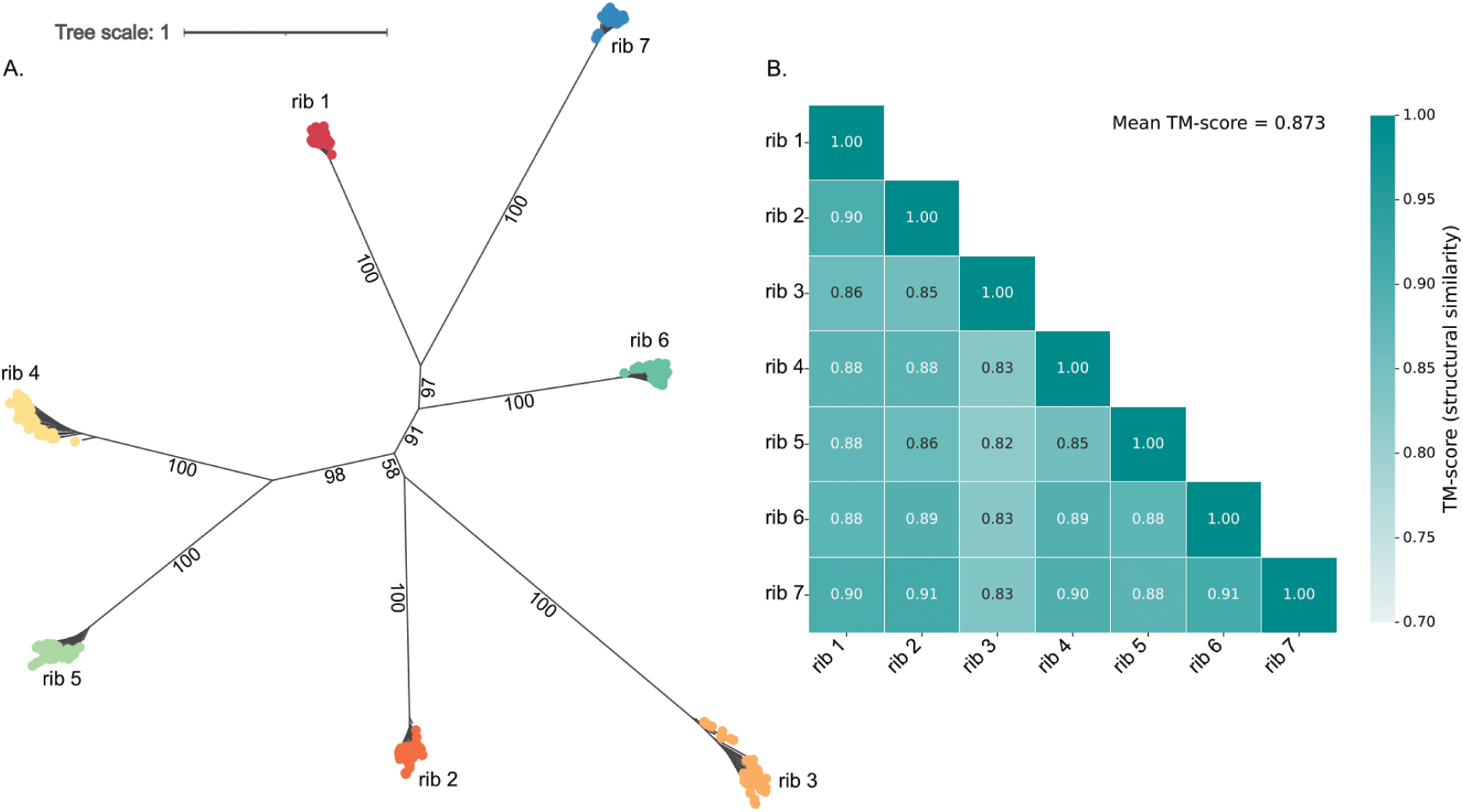
Phylogenetic analysis and structural conservation of *rib* modules in *T. cruzi* Dm28c. (A) Unrooted phylogenetic tree constructed from 448 *rib* modules (64 full-length DGF-1 copies with 7 modules per copy) identified in the *T. cruzi* Dm28c strain. The clustering patterns reflect the evolutionary relationships among the repetitive units composing DGF-1 proteins. (B) Structural similarity matrix (Heatmap) based on pairwise TM-score comparisons among individual *rib* modules derived from representative full-length architecture. Modules were modeled using AlphaFold3 and subjected to pairwise structural alignments.

To further evaluate structural relationships among *rib* modules, we performed all-against-all structural comparisons and calculated pairwise TM-scores (Figure 4B). *Rib* modules show high structural similarity, with TM-scores ranging from 0.82 to 0.91 (mean = 0.87), indicating a shared structural fold across all modules (Xu and Zhang, 2010).

Together, these results indicate that DGF-1 proteins are composed of a series of structurally conserved but sequence-divergent modules.

### 5. Evolution of the DGF-1 modular architecture

To investigate the evolutionary origin of the DGF-1 family, we performed domain-resolved homology searches across a broad collection of Discoba proteomes, with particular emphasis on kinetoplastids. Based on the AlphaFold3 structural models described above, we separately extracted the *rib* modules and the C-terminal region containing the *hat* domain and transmembrane helices. These sequences were used to construct domain-specific hidden Markov model (HMM) profiles (Alvarez-Carreño et al. 2026), which were subsequently employed to screen these proteomes.

Screening 86 Euglenozoa genomes, most of which were highly complete (mean BUSCO completeness = 94%; only six assemblies showed values below 70%; Supplementary Table S3), revealed that the complete DGF-1 architecture is restricted to a subset of trypanosomatids, whereas its individual structural modules have a considerably broader evolutionary distribution (Figure 5A; Supplementary Table S4). No significant matches were detected in non-kinetoplastid Discoba representatives, including *Naegleria gruberi*, *Andalucia godoyi*, *Euglena gracilis*, and *Diplonema papillatum*. In contrast, free-living bodonids such as *Bodo saltans* and *Neobodo designis* contained numerous proteins harboring either *rib*-like modules or the *hat* and transmembrane domains. However, no proteins containing these three structural domains within the same polypeptide chain were identified in these organisms.

**Figure 5.**
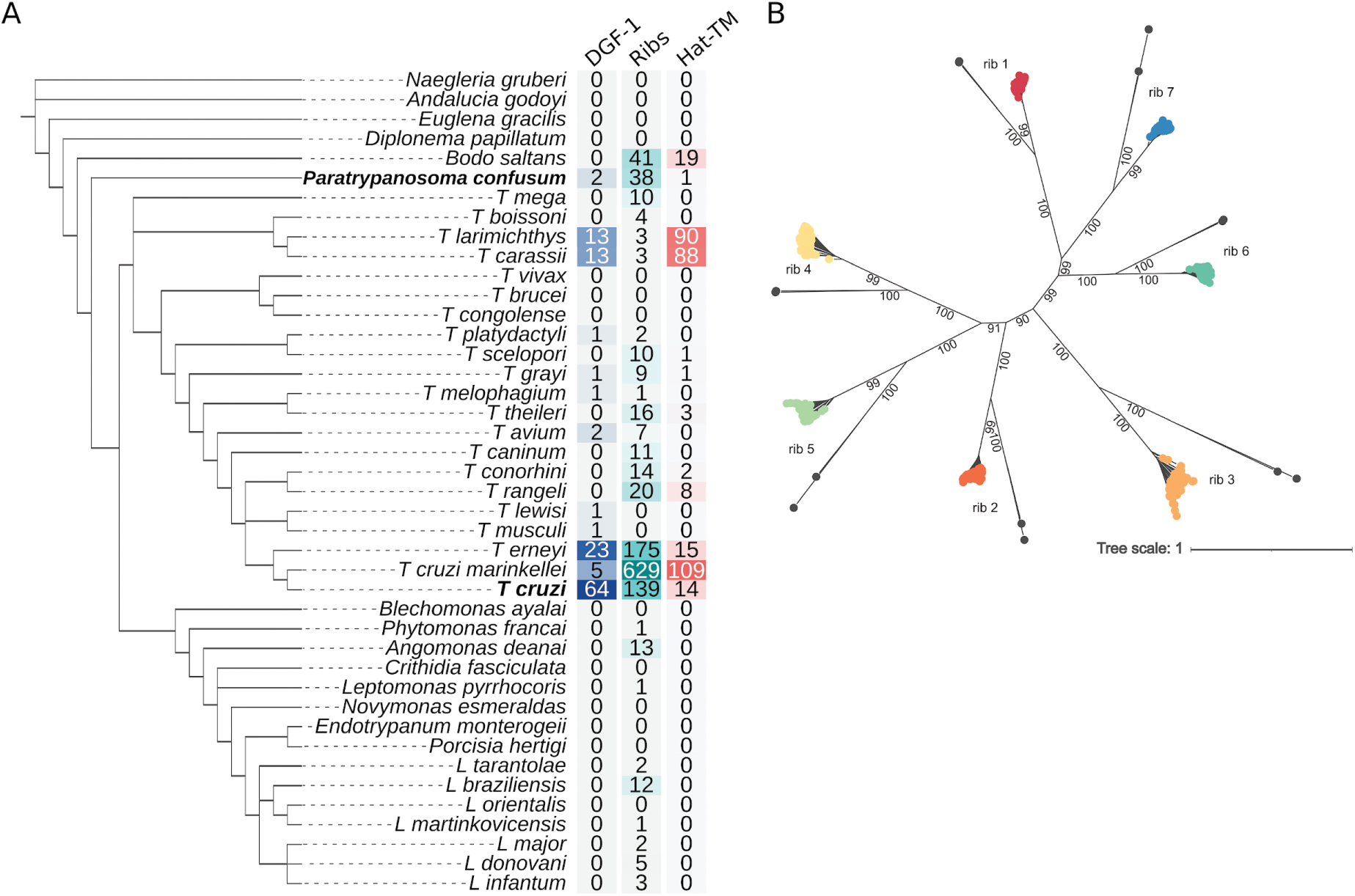
(A) Evolutionary distribution and lineage-specific expansion of the DGF-1 family across Discoba. Cladogram showing the evolutionary relationships among the Discoba species analyzed in this study, adapted from (Yang et al. 2026; Valach et al. 2023; Kostygov et al. 2024). The first column indicates the number of complete DGF-1 proteins identified in each genome, defined as proteins containing the complete canonical DGF-1 architecture. The second and third columns indicate the total number of *rib* domains and C-terminal *hat* domains associated with transmembrane helices (TM) identified in each genome. (B) **Phylogenetic analysis and structural conservation of *rib* modules between *T. cruzi* Dm28c and *P. confusum*.** Unrooted phylogenetic tree constructed from 462 *rib* modules, including 448 modules from *T. cruzi* Dm28c in colors (64 full-length copies with 7 modules per copy) and 14 modules from *P. confusum* in gray (2 full-length copies with 7 modules per copy).

We identified two full-length DGF-1 proteins in the *P. confusum* genome displaying the canonical seven-rib organization observed in *T. cruzi*, together with numerous isolated *rib*-like modules and independent transmembrane-associated domains. The phylogenetic position of *P. confusum* and the conservation of the complete DGF-1 architecture prompted a detailed comparison of its *rib* modules with those of *T. cruzi*.

Structural comparisons revealed remarkable conservation between *P. confusum* and *T. cruzi ribs*, with TM-scores ranging from 0.81 to 0.97 (mean = 0.87) (Supplementary Figure S10). Consistently, phylogenetic reconstruction placed the *P. confusum rib* sequences at the base of each of the seven positional *rib* clades identified in *T. cruzi* (Figure 5B). Together, these results indicate that the complete seven-rib DGF-1 architecture, as well as the positional identity of each *rib* module, had already evolved before the diversification of trypanosomatids.

Following the emergence of the complete DGF-1 architecture, it underwent different evolutionary pathways. Most *Trypanosoma* species retained only a small repertoire of complete genes. Non-salivarian trypanosomes, including *Trypanosoma grayi*, *T. rangeli*, *T. theileri*, and *T. avium*, typically contain one or two complete DGF-1 copies, suggesting that a low-copy repertoire represents the ancestral state in most trypanosomatids. Although genome completeness may influence copy number estimates in some species (Supplementary Table S3), this pattern was consistently observed across the majority of analyzed genomes.

Whereas, African trypanosomes, including *Trypanosoma brucei*, *T. vivax*, and *T. congolense*, lacked detectable DGF-1 genes or DGF-1 related domains, indicating a complete secondary loss of the family. On the other hand, in *Leishmania* and other non *Trypanosoma* species (e.g. *Blastochritidia*, *Leptomonas*, etc) complete DGF-1 proteins and transmembrane-associated domains were not detected, although several species retained isolated *rib*-like modules, suggesting a lineage-specific reduction of the ancestral DGF-1 architecture rather than complete elimination of its structural components.

*Trypanosoma cruzi* has the most extensive expansion of the DGF-1 family. Although only five complete DGF-1 genes, together with numerous isolated *rib* and *hat*–TM modules, were recovered from *T. cruzi marinkellei,* this likely reflects the fragmented nature of the available genome assembly rather than the true size of the family (Figure 5B, Supplementary Table S3). It is worth noting, however, that the expansion of the DGF-1 family is not unique to *T. cruzi.* Its close relative *T. erneyi* contains 23 complete DGF-1 genes, while *T. larimichthysi* and its very close relative *T. carassii* each harbor 13 complete copies. Consistent with the species distribution of gene copy numbers, phylogenetic reconstruction of all complete DGF-1 proteins revealed largely lineage-specific clustering supporting independent expansion (Supplementary Figure S12).

### 6. Evolutionary origins of DGF-1

In the previous section, we presented evidence indicating that DGF-1 was already present in the ancestor of all trypanosomatids, since the earliest branching *P. confusum* contains complete DGF-1 copies. Nevertheless, it is of interest to trace back the evolutionary emergence of this protein.

Our HMM search identified 41 *rib*-like and 19 *hat*-TM domains in *B. saltans*. Structural comparisons of *rib*-like domains show that these modules have the same right-handed parallel β-helix fold as DGF-1 *ribs*. Jackson et al. proposed that a subset of bodonins might represent the ancestral forms of DGF-1 proteins, but the representative bodonins highlighted in their study do not display either the characteristic tandem *rib* architecture that defines DGF-1 proteins or the *hat* domain (Supplementary Figure S8).

To further investigate the relationship between DGF-1 and *B. saltans*, we performed a structure-based homology search against the predicted structural proteome of *B. saltans* using the *T. cruzi* DGF-1 structure as the query (Supplementary Table S5). We identified 53 proteins with significant structural similarity to DGF-1. Of these, 21 were annotated as bodonins, all of which aligned exclusively to the C-terminal membrane-associated region, with no detectable similarity to the *rib* modules. Among the remaining 32 non-bodonin proteins, four matched *rib* modules, whereas the other 28 aligned to the C-terminal membrane-associated region.

Three additional proteins identified by the *rib*-specific HMM searches (BSAL_77525, BSAL_72995, and BSAL_27460) are of particular interest because they contain multiple *rib*-like modules (9, 4, and 3, respectively).

We reconstructed a structure-based phylogeny including all *rib*-like domains identified in *B. saltans*, together with the *rib* domains from *T. cruzi* and *P. confusum* (Supplementary Figure S9). All trypanosomatid *rib* domains, with the exception of *rib* 3, formed a well-supported clade composed exclusively of trypanosomatid sequences. In contrast, *rib* 3 clustered with the *B. saltans rib*-like domains.

Our findings are consistent with the proposed evolutionary relationship between the bodonin transmembrane domain and the corresponding region of DGF-1. Moreover, we found that this domain is not restricted to bodonin proteins. Importantly, our analyses show that bodonins lack the tandem *rib* architecture that characterizes DGF-1 proteins (Supplementary Figure S8). Instead, *rib*-like structures are present in *B. saltans* proteins outside the bodonin family, a finding that, to our knowledge, is reported here for the first time. Together, these observations suggest that proteins structurally related to both the *rib* and the *hat*–TM modules are present in *B. saltans*, although these modules were not found within the same protein.

## Discussion

Multigene families are a defining feature of the *Trypanosoma cruzi* genome and have traditionally been associated with host interaction, immune evasion, and parasite adaptation. Among them, DGF-1 represents one of the largest genomic investments made by the parasite, yet its biological role has remained remarkably elusive. Despite comprising hundreds of copies encoding unusually large proteins, DGF-1 has received considerably less attention than other expanded families such as trans-sialidases or mucins. Experimental evidence indicating that DGF-1 proteins are predominantly expressed in intracellular amastigotes rather than infective trypomastigotes (Lander et al. 2010) suggests that their function may differ from that of classical surface antigens and instead reflect a more specialized role during intracellular development.

In this study, we combined curated gene annotation, structural prediction, and comparative genomics to shed some light on the evolutionary history of the DGF-1 family. A central finding of our work is the identification of seven tandemly arranged structural modules, here termed *ribs*, that constitute the extracellular region of DGF-1 proteins. Each rib adopts a right-handed parallel β-helix fold. Together, these modules, followed by a conserved C-terminal *hat*–TM membrane-associated region, define a previously unrecognized modular architecture. This structural framework provided a level of evolutionary resolution that could not be achieved using sequence similarity alone. In particular, the identification of individual *rib* modules enabled the construction of domain-specific HMM profiles capable of detecting homologous structural elements across deeply divergent kinetoplastids, substantially extending previous evolutionary analyses of the family (Jackson et al. 2016; Kawashita et al. 2009a; Ramírez 2023).

Our comparative analyses revisit the evolutionary proposal made by Jackson et al. 2016. Although bodonins have been suggested as the ancestral lineage of DGF-1 proteins, our structural analyses indicate that bodonin proteins do not represent the direct ancestral form of DGF-1, as they lack the defining architecture of this family. Instead, our results suggest that the *rib* domain represents the ancestral structural module that was subsequently incorporated into the canonical DGF-1 architecture. In contrast, the earliest branching trypanosomatid, *P. confusum* (Flegontov et al. 2013), already possesses complete DGF-1 proteins displaying the same overall architecture observed in *T. cruzi*. Together, these observations strongly support a model in which the defining evolutionary innovation was the assembly of the complete multidomain DGF-1 architecture early during trypanosomatid evolution, whereas the individual structural components have arisen before the origin of the family.

Equally remarkable is the degree of architectural conservation maintained throughout the diversification of trypanosomatids. The seven *rib* modules are not simple tandem repeats but evolutionarily differentiated structural units. Across all analyzed DGF-1 copies, each *rib* consistently clusters with its positional counterpart rather than with neighboring modules within the same protein, indicating that the identity and relative order of the seven modules have been preserved since the emergence of the family. The observation that *P. confusum*, retains the same modular organization as *T. cruzi* suggests that this architecture has been maintained under strong evolutionary constraint since the origin of the *trypansomatid* family. Such conservation strongly supports the idea that individual *ribs* contribute with distinct structural or functional properties to the mature protein.

The evolutionary history of DGF-1 also reveals a clear distinction between the origin of the protein architecture and the subsequent expansion of the gene family. Most *Trypanosoma* species retain only one or a few complete DGF-1 genes, indicating that the multidomain organization remained evolutionarily stable long before extensive gene amplification occurred. Large repertoires appear to have evolved only in a limited number of lineages, most prominently within the *Schizotrypanum* clade, although an independent expansion is also observed in *T. larimichthysi* and *T. carassii*. Notably, high-quality genome assemblies for these latter species have only recently become available, highlighting the importance of continued genome sequencing across trypanosomatids (Liu et al. 2026; Lv et al. 2026). As additional high-quality genomes become available, the evolutionary distribution of DGF-1 proteins will likely be further refined, and additional lineage-specific expansions may yet be identified.

Conversely, African salivarian trypanosomes have completely lost both DGF-1 and its constituent structural modules, whereas *Leishmania* and other no-*Trypanosoma* trypanosomatids species retain only isolated *rib*-like domains. Together, these patterns suggest that the complete DGF-1 architecture emerged once early during trypanosomatid evolution and subsequently experienced lineage-specific amplification, reduction, and several secondary losses. The complete loss of both DGF-1 and its ancestral *rib* modules in African trypanosomes further indicates that, despite the deep evolutionary conservation of the family architecture, its maintenance is not universally required across trypanosomatids and likely depends on lineage-specific ecological or host-associated selective pressures. This evolutionary pattern resembles that described for other trypanosomatid surface protein families, such as GP63, which exhibit an ancient origin followed by lineage-specific expansions across different evolutionary lineages (Berná et al. 2025). Together, these observations suggest that the emergence of a conserved protein architecture and some subsequent expansions of the corresponding gene family represent distinct evolutionary processes that have shaped the diversification of trypanosomatid surface proteins.

The identification of the *rib* modules also provides new perspectives on the possible function of DGF-1 proteins. Right-handed parallel β-helices constitute one of the best-characterized structural scaffolds in extracellular proteins. Most functionally characterized members of this fold are enzymes involved in the degradation of complex polysaccharides, although the same structural scaffold has also been adapted for roles in adhesion and macromolecular recognition (Jenkins y Pickersgill 2001). Although InterProScan assigns several DGF-1 regions to pectin lyase-like domains, these annotations likely reflect recognition of a shared right-handed parallel β-helix scaffold rather than conserved enzymatic function. Likewise, the predicted EGF-like and lipid-binding domains probably result from local structural similarities rather than true homology.

These findings also illustrate the benefits of combining profile-based annotation methods, with protein structure approaches. Our results therefore emphasize the value of structure-guided analyses for reconstructing evolutionary relationships that cannot be resolved by sequence-based annotation alone (Al-Fatlawi et al. 2023).

Another unexpected finding is the presence of the previously undescribed *hat* domain linking the *ribs* region to the transmembrane helices. Structural comparisons consistently classify this domain within the immunoglobulin-like β-sandwich superfamily despite the complete absence of detectable sequence similarity. Although its precise biological role remains unknown, its conserved position immediately preceding the membrane anchor suggests that it may act as a structural interface between the extracellular *rib* modules and the membrane-associated region. Whether this domain contributes to protein stability, oligomerization, or interactions with parasite or host molecules remains an important question for future functional studies.

Finally, our work illustrates the power of integrating comparative genomics with protein structure prediction to resolve the evolutionary history of highly repetitive gene families. Sequence similarity alone was insufficient to recognize the modular organization of DGF-1 or to reconstruct its evolutionary origin across deeply divergent kinetoplastids. By incorporating structural information, we were able to identify the ancestral building blocks of the family, reconstruct the emergence of its multidomain architecture, and distinguish between the origin of the protein itself and the later lineage-specific expansions that produced the remarkable DGF-1 repertoires observed in *T. cruzi*. We propose that DGF-1 represents an ancient modular innovation that originated early during trypanosomatid evolution through the assembly of pre-existing structural elements and whose exceptional expansion in *T. cruzi* reflects a secondary evolutionary process acting on an already established and highly conserved protein architecture.

## Materials and methods

### Genome assemblies and proteome datasets

In this study, we used the telomere-to-telomere (T2T) genome assembly of *Trypanosoma cruzi* Dm28c generated from PacBio HiFi sequencing data, comprising 32 chromosomes with fully separated haplotypes (Greif et al. 2026). Genome assemblies and predicted proteomes from 91 additional representative Discoba species were retrieved from GenBank (Benson et al. 2014), TriTrypDB (Amos et al. 2022), EukProt (Richter et al. 2022), and other publicly available repositories (Supplementary Table S4). As the initial reference set for DGF-1 identification, we used the annotation of DGF-1 genes previously generated for the Dm28c strain by Berná et al. (2018).

### Genome quality assessment

Genome completeness was assessed for all analyzed genome assemblies using BUSCO v5.8.3(Manni et al. 2021) with the *euglenozoa_odb10* lineage dataset under default parameters. Most genomes showed high completeness, with a mean BUSCO score of approximately 94%, whereas six genomes displayed completeness values below 70%. BUSCO completeness metrics for all analyzed genomes are provided in Supplementary Table S3 and Supplementary Figure S11.

### Identification and re-annotation of DGF-1 genes and pseudogenes

DGF-1 loci were identified using a pipeline combining BLASTn v2.12.0+, BLASTp v2.12.0+, (Altschul et al. 1990) transeq, and getorf from the EMBOSS package v6.6.0.0 (Olson 2002), together with custom R scripts developed for this study (available at: https://github.com/mathiashole/Blast-and-HMM-searches). Searches were performed using BLAST with default parameters unless otherwise specified.

The initial query set consisted of a manually curated collection of DGF-1 sequences from the *T. cruzi* Dm28c strain described by (Berná et al. 2018), available at http://bioinformatica.fcien.edu.uy/cruzi. This reference set was used for homology-based identification and validation of DGF-1 loci.

To define gene boundaries, fragmented BLASTn hits separated by less than 100 bp were merged, allowing the reconstruction of disrupted loci and fragmented gene models derived from the same ancestral sequence. Candidate sequences were classified as intact DGF-1 genes if they met the following criteria: (i) at least 80% coverage relative to the expected full-length DGF-1 protein (∼3,000 amino acids), (ii) a minimum sequence identity of 80% relative to canonical DGF-1 members, and (iii) the presence of a continuous open reading frame (ORF). These thresholds were selected based on the expected size and conservation of canonical DGF-1 genes.

Sequences failing to meet these criteria were considered putative pseudogenes and subjected to further analysis. Pseudogene annotation was based on a combination of sequence similarity and coding potential. Genomic regions showing homology to DGF-1 genes but lacking a continuous ORF, containing ORFs shorter than 80% of the expected protein length, or displaying reduced sequence identity (<80%) were classified as pseudogenes. To confirm their relationship to the DGF-1 family, all pseudogene candidates were required to retain detectable homology to canonical DGF-1 proteins, defined as a minimum e-value ≤ 1e−5.

Together, these criteria allowed the identification of a high-confidence set of intact DGF-1 genes and pseudogenes while excluding unrelated sequences.

### Analysis of chromosomal distribution and sequence composition

The chromosomal distribution of DGF-1 genes and pseudogenes was analyzed by mapping their genomic coordinates across the assembly. To account for differences in chromosome size, locus positions were normalized as a fraction of total chromosome length. Visualizations of gene distribution and positional density were generated using custom R scripts (available at: https://github.com/mathiashole/chromR).

GC content was calculated for all DGF-1 genes, pseudogenes, and annotated protein-coding genes from the Dm28c genome using infoseq with custom bash script. Trinucleotide composition analyses were performed using normalized trinucleotide frequencies calculated for each sequence. Principal component analysis (PCA) based on trinucleotide composition was performed in R version 4.5.0 using the prcomp function under default settings.

Differences between observed locus distributions and uniform chromosomal distributions were evaluated using Wilcoxon rank-sum tests implemented in R.

### Domain annotation and sequence analyses

Conserved domains in DGF-1 proteins were identified using InterProScan v5.76-107.0 with default parameters. InterPro annotations were parsed and summarized using custom R scripts to reconstruct domain architectures across the family available at: https://github.com/mathiashole/ipegg.

Multiple sequence alignments of DGF-1 genes were generated separately for nucleotide and amino acid sequences using MAFFT v7.526 with the --auto option. Pairwise sequence identity values were computed from the resulting alignments using a custom Python script based on Biopython, considering only alignment positions without gaps in either sequence.

Identity matrices were converted into distance matrices (1 − identity) and used to generate clustered heatmaps. Hierarchical clustering was performed using average linkage and Euclidean distance as implemented in the seaborn Python library v.3.8.10

### Protein structure prediction and structural analyses

Protein structure predictions were generated using AlphaFold3 for representative DGF-1 sequences from different *T. cruzi* phylogenetic groups, as well as for inferred ancestral homologs identified in related kinetoplastids. In addition to full-length proteins, individual *rib* modules and the C-terminal transmembrane-associated region were modeled independently.

Predicted structures were visualized and analyzed using UCSF ChimeraX v1.12 (Pettersen et al. 2021)and Mol* v5.8.0 Viewer (Rose et al. 2026). Structural similarity among *rib* domains was evaluated using Foldseek release 10-941cd33 (van Kempen et al. 2024) through all-against-all structural comparisons. Pairwise TM-scores were used as quantitative measures of structural similarity among domains.

To identify structural homologs and classify DGF-1 domains, predicted structures were queried against the CATH database (Waman et al. 2025) using the Foldseek web server. Structural matches were filtered using a TM-score threshold > 0.5 and an e-value <1e−10, with all structural similarity metrics (TM-score) corresponding to the structural superposition recalculated by the Foldseek web viewer interface for each selected hit.

### Structural similarity search for DGF-1 homologs in *B. saltans*

A structural similarity search was performed to identify remote homologs of DGF-1 in *Bodo saltans*. The complete set of predicted protein structures available for *B. saltans* was downloaded from the AlphaFold Protein Structure Database (Jumper et al. 2021; Bertoni et al. 2026) and used as the basis for the target database (18.139 protein structures). To obtain a comprehensive structural dataset, additional proteins absent from the AlphaFold database were incorporated. These included 3 bodonin and 7 proteins identified by our *rib* HMM search. The resulting collection constituted the complete target structural database.

The predicted structure of *T. cruzi* DGF-1 was used as the query. Structural similarity searches were performed using Foldseek (10.941cd33 version) with the following parameters: --exhaustive-search 1 --exact-tmscore 1. Hits were retained using as filtering criteria of TM-score > 0.5 and E-value < 1 × 10−5, resulting in 54 protein structure hits.

Candidate homologs were further examined in the context of orthologous groups obtained from OrthoMCL clusters available in TriTrypDB release 70. Proteins were classified as bodonins if they belonged to the same OrthoMCL groups as BSAL_90090, BSAL_52525, BSAL_08390, BSAL_73585, BSAL_90835, BSAL_11510, BSAL_31875, BSAL_37140, BSAL_11320, BSAL_92780 based on (Jackson et al. 2016). Resulting in 388 bodonin proteins distributed in the OrthoMCL groups: OG8_0011510, OG8_0008752, OG8_0007749, OG8_0003662, OG8_0039660, OG8_0059364, OG8_0016555, OG8_0017661. BSAL_11510, was not represented in TriTrypDB so no OrthoMCL group has been assigned.

### Phylogenetic analyses

Phylogenetic relationships were reconstructed using five independent datasets: (i) full-length DGF-1 proteins from *T. cruzi* Dm28c haplotype 1; (ii) full-length DGF-1 proteins from multiple *T. cruzi* strains (Dm28c, TCC, YC6, and Dm25 haplotype 1); (iii) individual *rib* modules from *T. cruzi* Dm28c, with each module treated as an independent sequence; (iv) *rib* modules including homologous sequences from *Paratrypanosoma confusum*; and (v) full-length DGF-1 homologs identified across kinetoplastids. Trees corresponding to datasets (i) and (ii) were rooted using representative DGF-1 homologs from *Trypanosoma erneyi* as an outgroup.

Multiple sequence alignments were generated using MAFFT v7.526 with up to 1,000 iterative refinements (Katoh y Standley 2013). Poorly aligned regions were automatically trimmed using trimAl v1.5.rev0 with the -automated1 heuristic option.

Maximum-likelihood phylogenetic trees were reconstructed using IQ-TREE v2.0.7 (Wong et al. 2026). Best-fit amino acid substitution models were independently selected for each dataset using ModelFinder under the Bayesian Information Criterion (BIC). Branch support was evaluated using the Ultrafast Bootstrap 2 (UFBoot2) approximation with 1000 replicates.

In addition to sequence-based phylogenetic analysis, structural relationships among DGF-1 *rib* domains from *T. cruzi*, *P. confusum*, and *rib*-like domains identified in *B. saltans* were reconstructed using FoldTree as implemented in Foldseek (Moi et al. 2025). FoldTree infers phylogenetic relationships directly from pairwise protein structural similarities without requiring multiple sequence alignment.

All phylogenetic trees were visualized and annotated using iTOL version 7.6 (Letunic y Bork 2024).

### Identification of ancestral DGF-1 homologs across Discoba

To investigate the evolutionary origin of DGF-1 proteins, profile hidden Markov models (HMMs) were generated for the two major structural components of the protein. Based on AlphaFold3 structural predictions, DGF-1 proteins were partitioned into their individual *rib* domains and the C-terminal transmembrane-associated *hat* domain. All *rib* domains were combined into a single multiple sequence alignment using MAFFT v7.526, and the resulting alignment was used to build a profile HMM with hmmbuild from the HMMER v3.3.2 package (Alvarez-Carreño et al. 2026). A second profile HMM was generated from an alignment of the C-terminal transmembrane-associated *hat* domains.

Multiple sequence alignments for each module were generated using MAFFT v7.526, and profile HMMs were built using hmmbuild from the HMMER v3.3.2 package. The *rib*-domain and transmembrane-region HMMs were used independently to search representative Euglenozoa and Discoba proteomes using hmmsearch. Domain-level matches were extracted from the --domtblout output, retaining significant hits with e-values ≤ 1e-5. For genomes lacking annotated proteomes, all open reading frames (ORFs) longer than 100 amino acids were extracted using a custom bash script (available at: https://github.com/mathiashole/Blast-and-HMM-searches) to generate protein datasets for HMM screening.

Candidate proteins were reconstructed according to the combination of detected domains and classified as: (i) complete DGF-1 architectures comprising seven tandem *rib* modules and the C-terminal transmembrane-associated region; (ii) proteins containing only *rib*-like modules; or (iii) proteins containing only the transmembrane-associated region.

Structural similarity between ancestral *rib*-like modules and canonical *T. cruzi* DGF-1 *rib* domains was evaluated using Foldseek structural alignments and TM-score calculations.

## Acknowledgements

M.J.M. was supported by a postgraduate scholarship from the *Agencia Nacional de Investigación e Innovación* (ANII, Uruguay; grant number POS_NAC_2024_1_183254). F.A.-V., A.P., J.T., and L.B. are members of the *Sistema Nacional de Investigadores* (SNI, ANII, Uruguay) and researchers of the *Programa de Desarrollo de las Ciencias Básicas* (PEDECIBA, Uruguay).

