## Supplementary information for "The modular origin and evolutionary expansion of the enigmatic DGF-1 protein family in trypanosomatids"

### Figure S1

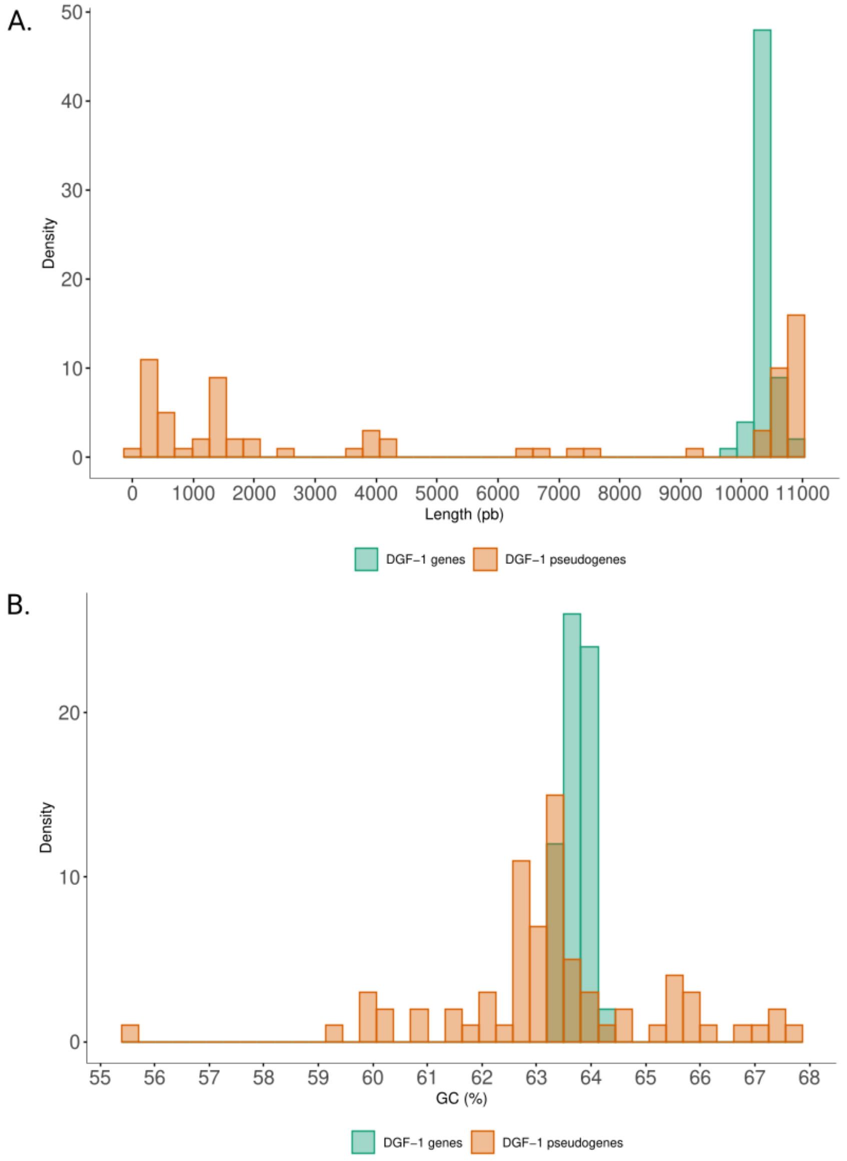


**Supplementary Figure S1. Distribution of DGF-1 sequence lengths and GC content in *T. cruzi* Dm28c.**

(A) Histogram showing the density distribution of DGF-1 sequence lengths, distinguishing intact genes (green) and pseudogenes (orange). (B) Histogram showing the density distribution of GC content in DGF-1 sequences, comparing intact genes (green) and pseudogenes (orange).

### Figure S2

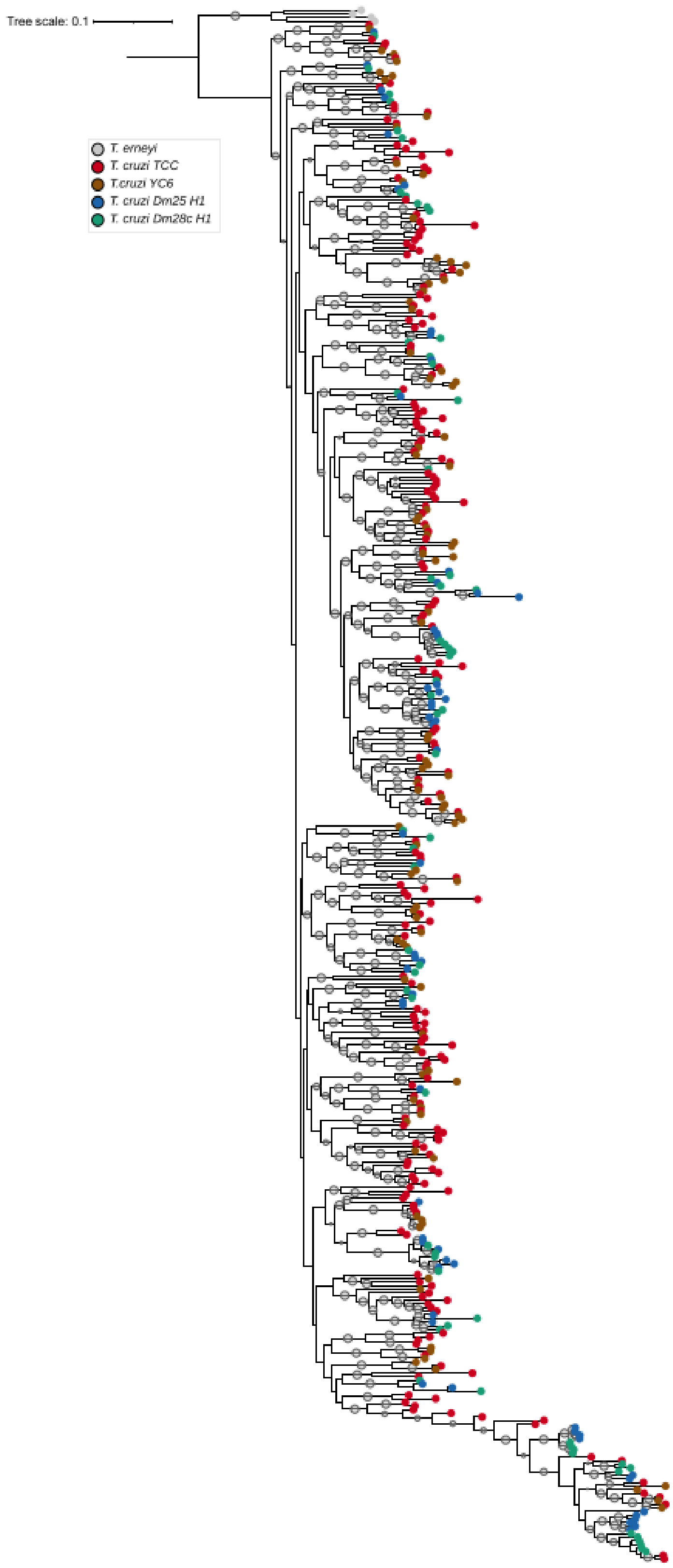

Tree scale: 0.1
T. erneyi
T. cruzi TCC
T.cruzi YC6
T. cruzi Dm25 H1
T. cruzi Dm28c H1

**Supplementary Figure S2. Phylogenetic relationships of the DGF-1 family from different *T. cruzi* strains.**

Rooted maximum-likelihood phylogeny reconstructed from full-length DGF-1 amino acid sequences identified in multiple *Trypanosoma cruzi* strains (Dm28c H1, Dm25 H1, TCC y YC6). The tree was rooted using DGF-1 sequences from *Trypanosoma erneyi* as the outgroup (indicated by gray circles at the branch tips). Only branch support values (≥ 95 UFBoot) are shown at nodes. Terminal branches are color-coded by strain according to the legend.

### Figure S3

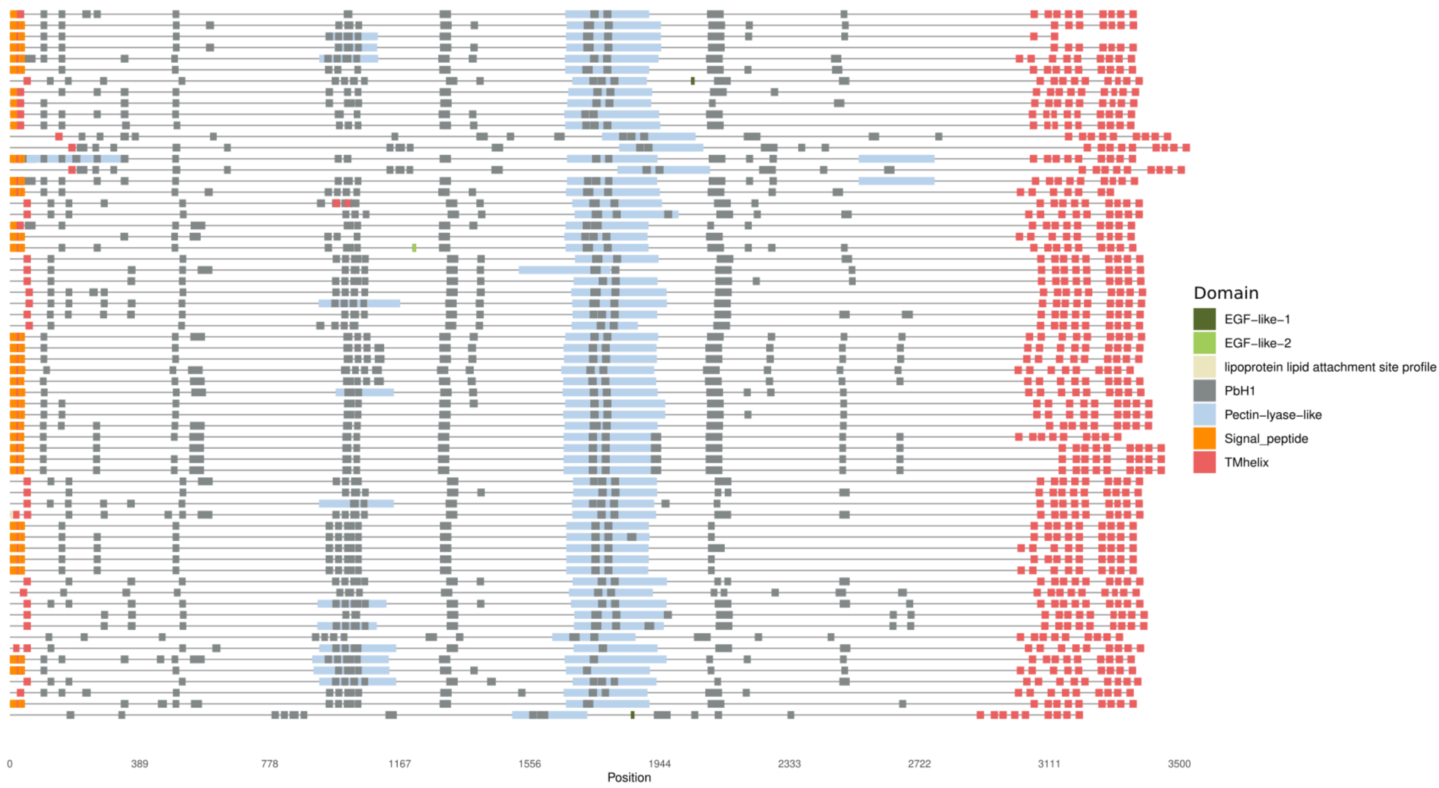


**Supplementary Figure S3. Comparative domain architecture of the DGF-1 multigene family.** Representative maps illustrating the diversity of domain organization across DGF-1 genes. Domain annotation was performed using InterProScan, and labels were standardized for clarity. Signal peptide (orange) and TMhelix (salmon) correspond to hydrophobic α-helical regions. In several genes, these features are annotated either as standalone TM helices or overlapping with signal peptides, reflecting their shared structural properties. Additional domains identified include: EGF-like-1 (dark green, PS00022); EGF-like-2 (light green, PS01186); PbH1 (gray, parallel β-helix repeats, SM00710); pectin lyase-like (light blue, SSF51126); and lipoprotein lipid attachment site (beige, PS51257). For visualization purposes, redundant annotations and low-confidence regions (including specific signal peptide subregions and selected Pfam entries such as PF11024 and PF11038) were removed. Genes are ordered alphanumerically.

### Figure S4

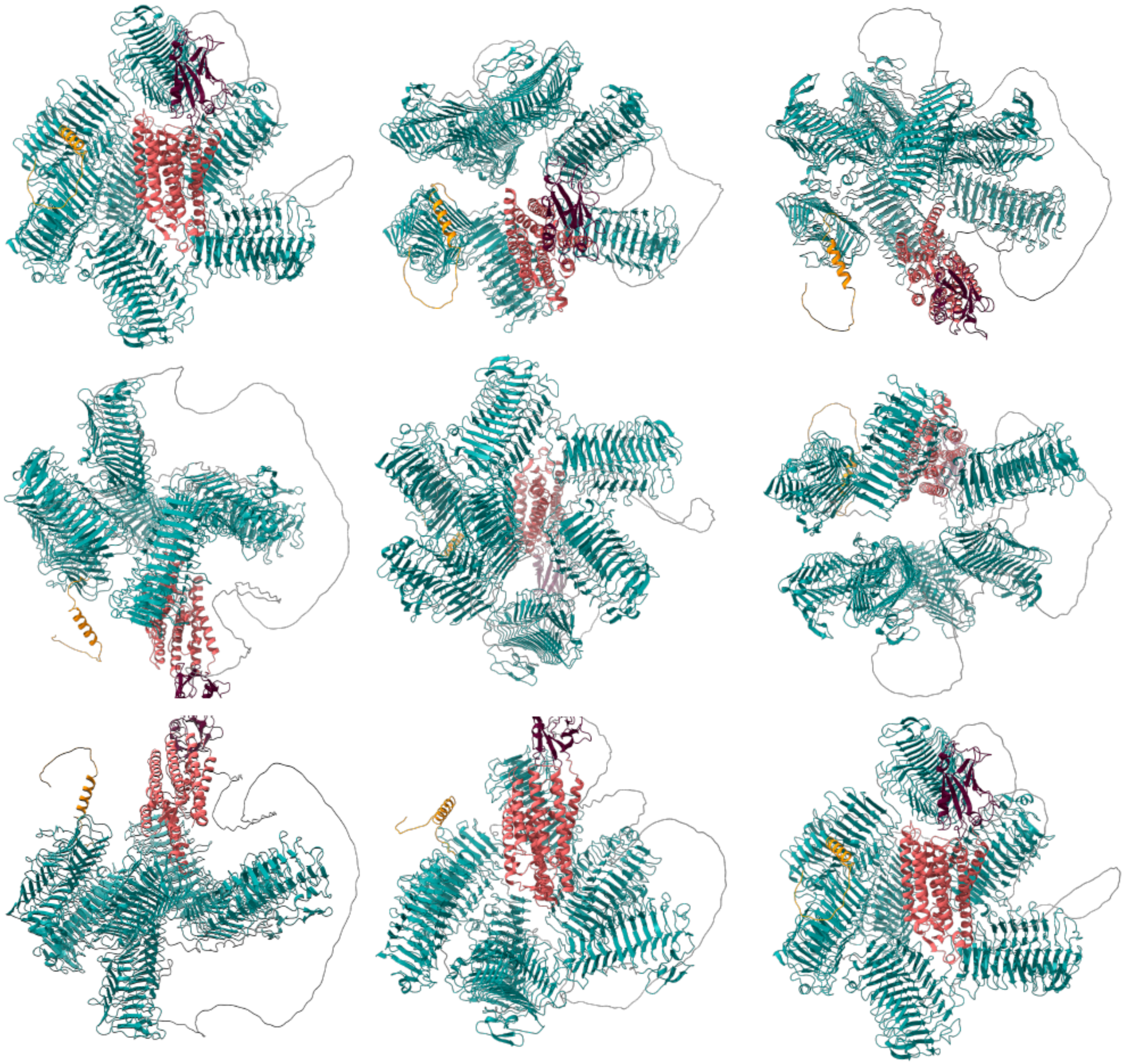

**Supplementary Figure S4. Three-dimensional structural prediction model of DGF-1.**

Full-length DGF-1 protein model predicted with AlphaFold3 and visualized using UCSF ChimeraX. The ribbon representation highlights the spatial arrangement of the signal peptide (orange), rib modules (teal), hat domain (wine), and transmembrane helices (salmon). Multiple orientations are shown to illustrate the overall fold and relative domain organization.

Figure S5

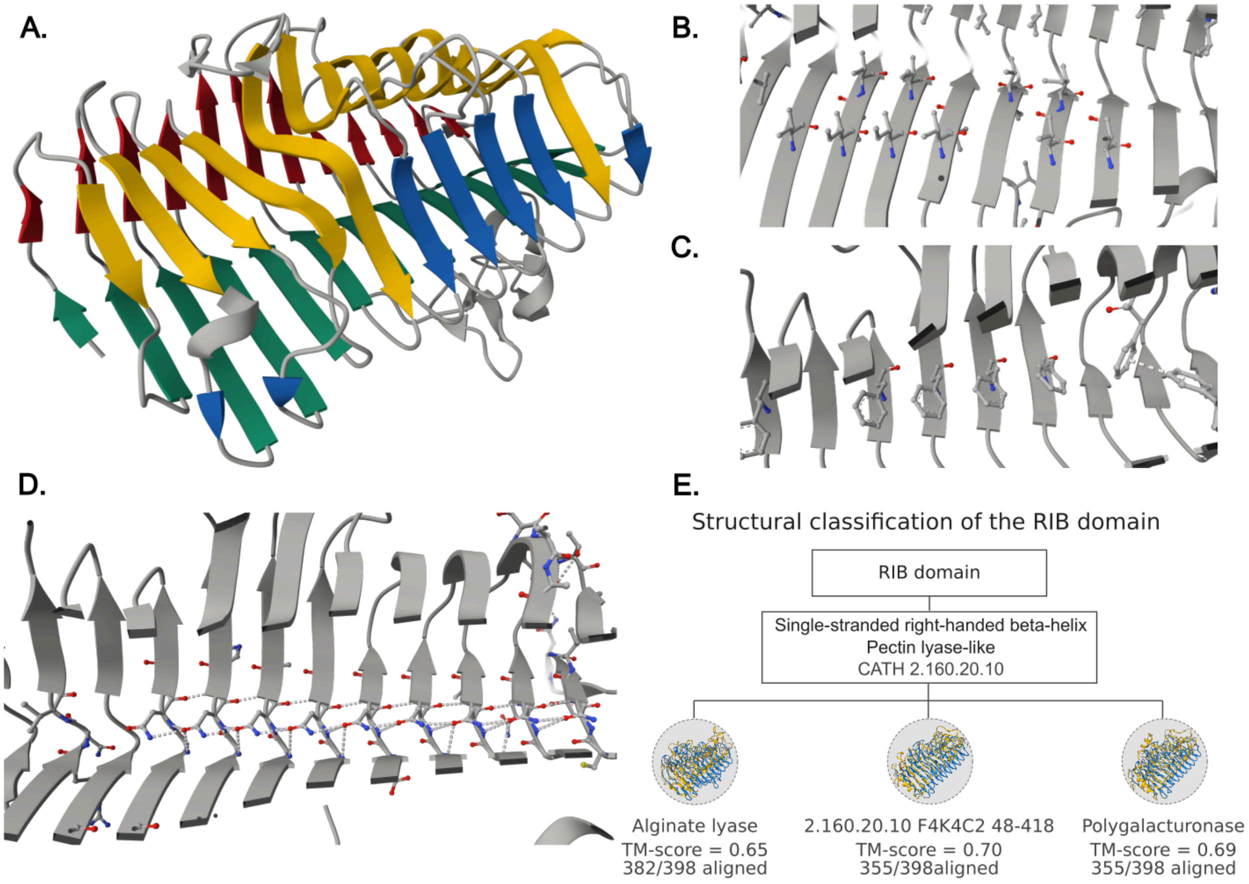


**Supplementary Figure S5. Structural architecture and characteristic stabilizing interactions of the DGF-1 Rib 5 module.**

(A) Ribbon diagram of a representative DGF-1 Rib 5 module illustrating the canonical right-handed parallel β-helix fold. The three principal β-sheets are colored according to the conventional nomenclature: PB1 (green), PB2 (yellow), and PB3 (red). The additional β-strands extending the PB1a face are shown in blue. (B) Representative aliphatic "cupped" stack formed by interdigitated isoleucine and valine side chains within the hydrophobic core , located between the PB1 and PB2 β-sheets. (C) Representative internal aromatic stack composed of phenylalanine residues arranged in an offset stacking configuration within the β-helix core along the PB3 β-sheets. (D) Internal hydrogen-bonding network forming a continuous asparagine ladder within the solenoid core. Dashed lines indicate hydrogen bonds between consecutive coils. Carbons are drawn in light grey with oxygens in red and nitrogens in blue. (E) Structural evidence supporting classification of the Rib domain as a right-handed parallel β-helix, Pectin lyase-like fold. The Rib domain was assigned to Single-stranded right-handed beta-helix, Pectin lyase-like folds (Superfamily 2.160.20.10) by Foldseek are shown together with their structural superpositions, TM-score and number of aligned residues. Structural renderings were generated using Mol*.

Figure S6

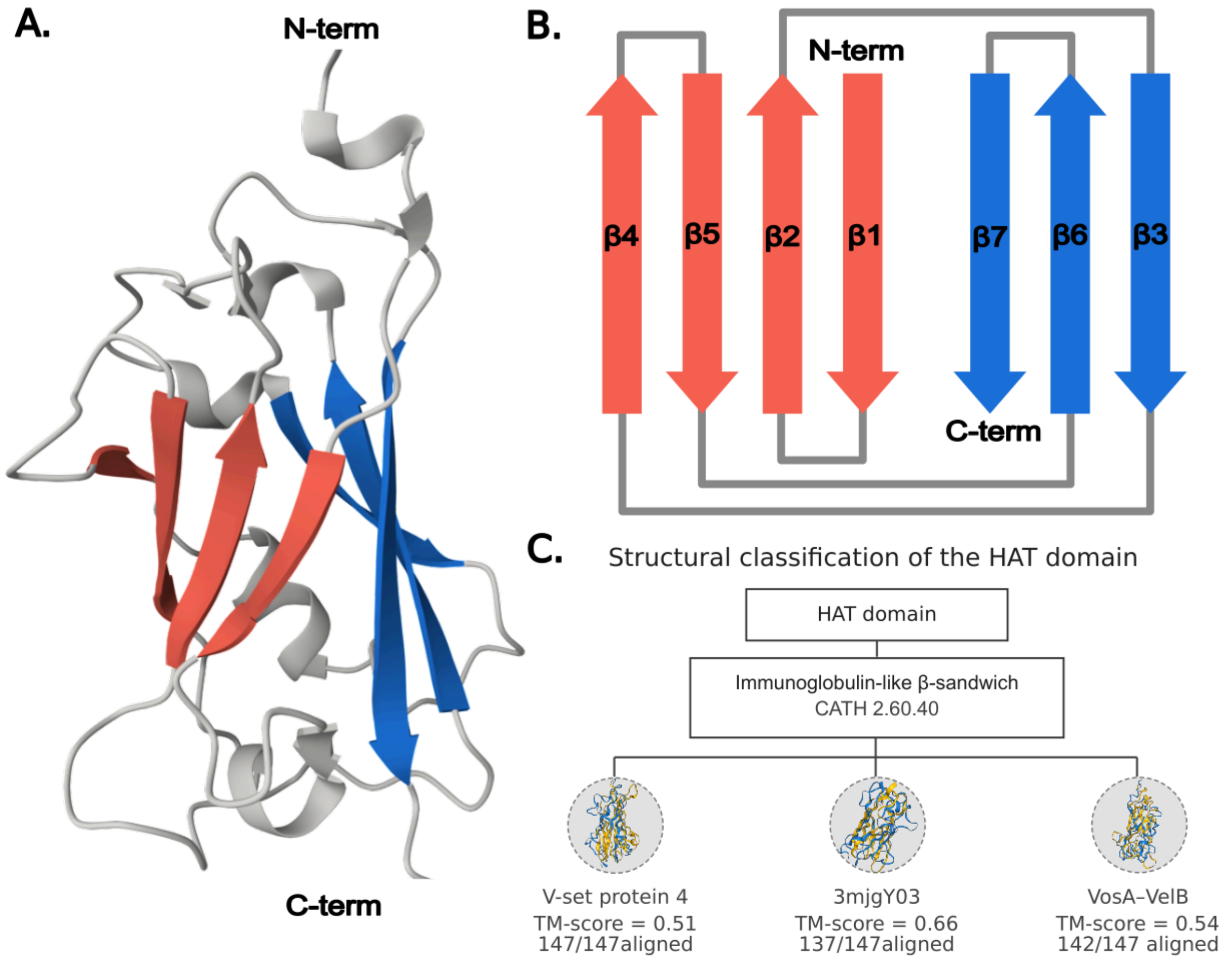


**Supplementary Figure S6. Structural characterization and classification of the Hat domain.**

(A) AlphaFold3-predicted structure of the Hat domain. The two opposing β-sheets forming the β-sandwich are highlighted in red and blue, whereas connecting loops and other secondary elements are shown in gray. N- and C-terminals are indicated. (B) Topology diagram of the Hat domain showing the connectivity and orientation of the seven β-strands. β-strands are colored according to the β-sheet to which they belong and arrow direction indicates the antiparallel arrangement of adjacent strands within the immunoglobulin-like β-sandwich fold. (C) Structural evidence supporting classification of the Hat domain as an immunoglobulin-like β-sandwich fold. The Hat domain was assigned to immunoglobulin-like β-sandwich fold (2.60.40.10), and representative structural matches belonging to CATH homologous superfamily 2.60.40.10 identified by Foldseek are shown together with their structural superpositions, TM-score and number of aligned residues.

### Figure S7

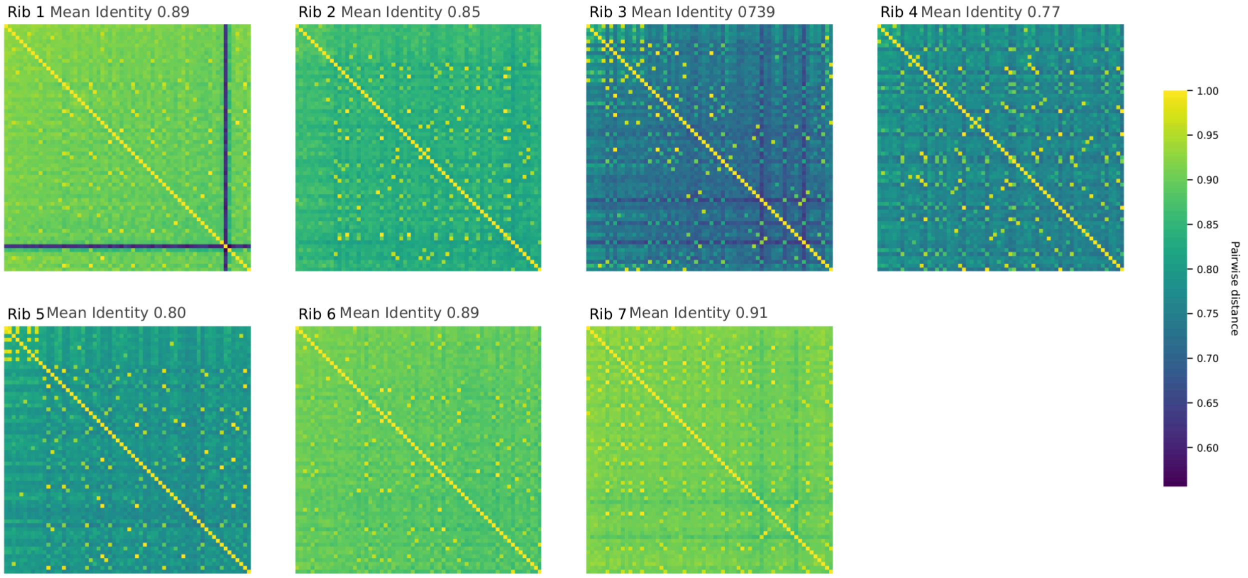


**Supplementary Figure S7. Pairwise sequence identity comparison across DGF-1 repeat units (Rib 1-7) in *Trypanosoma cruzi* Dm28c.**

Heatmaps displaying all-versus-all pairwise amino acid sequence identity matrices for each rib module (Rib 1 through Rib 7) across DGF-1 family members annotated in *Trypanosoma cruzi* Dm28c genome. The average pairwise identity ("Mean identity") for each specific repeat module is reported above its respective panel. The color gradient (viridis scale) indicates the degree of pairwise sequence identity, ranging from dark blue (low identity 0.50) to bright yellow (maximum identity 1.00).

### Figure S8

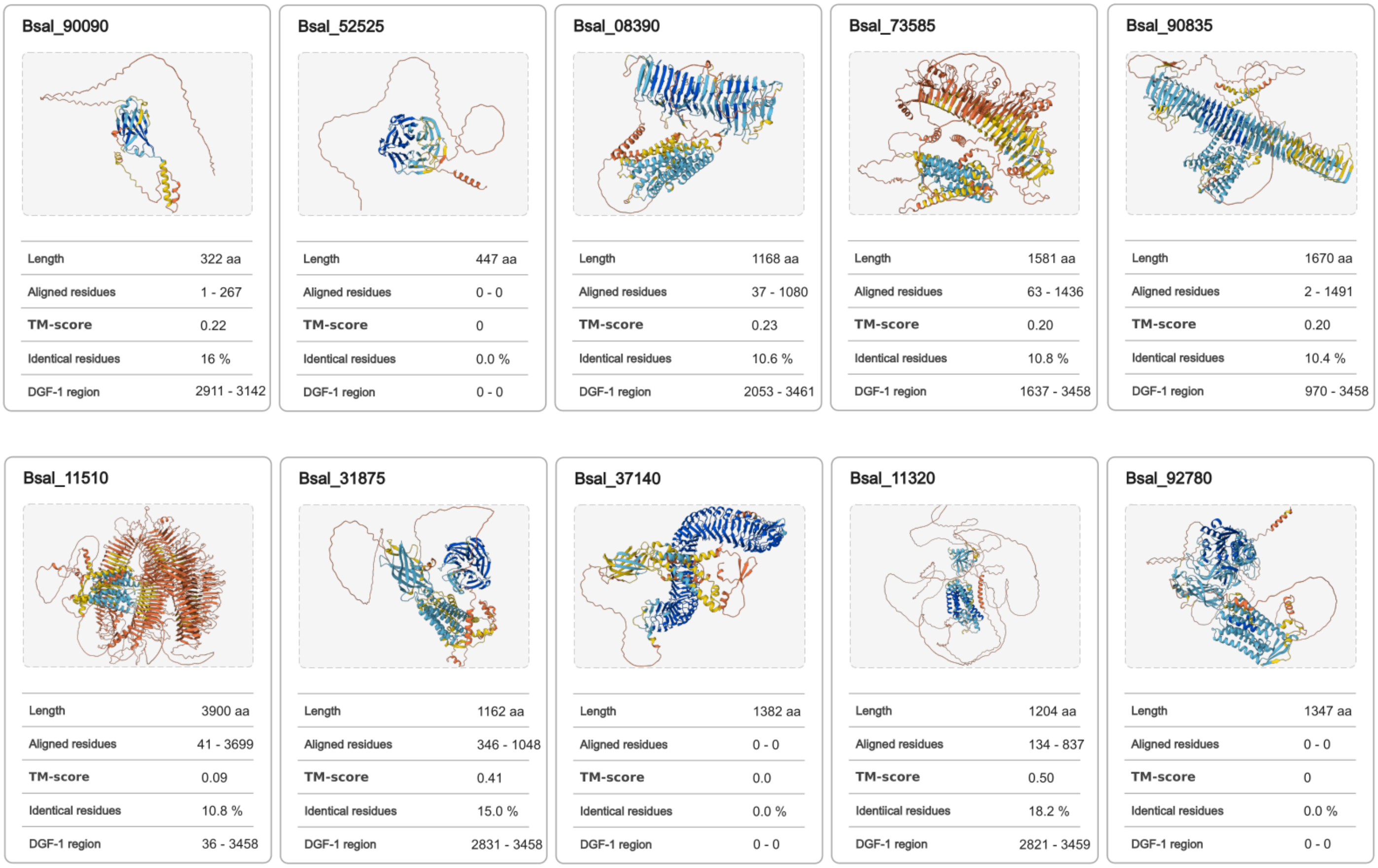


**Supplementary Figure S8. Structural comparison of *Bodo saltans* Bodonin isoforms with the *Trypanosoma cruzi* DGF-1 protein.**

AlphaFold3-predicted structures of the ten reported Bodonin isoforms were compared against the *Trypanosoma cruzi* DGF-1 protein using Foldseek. Each card summarizes the predicted structure and the corresponding structural comparison, including protein length, aligned regions in Bodonin and DGF-1 proteins, TM-score and sequence identity within structural alignment.

Figure S9

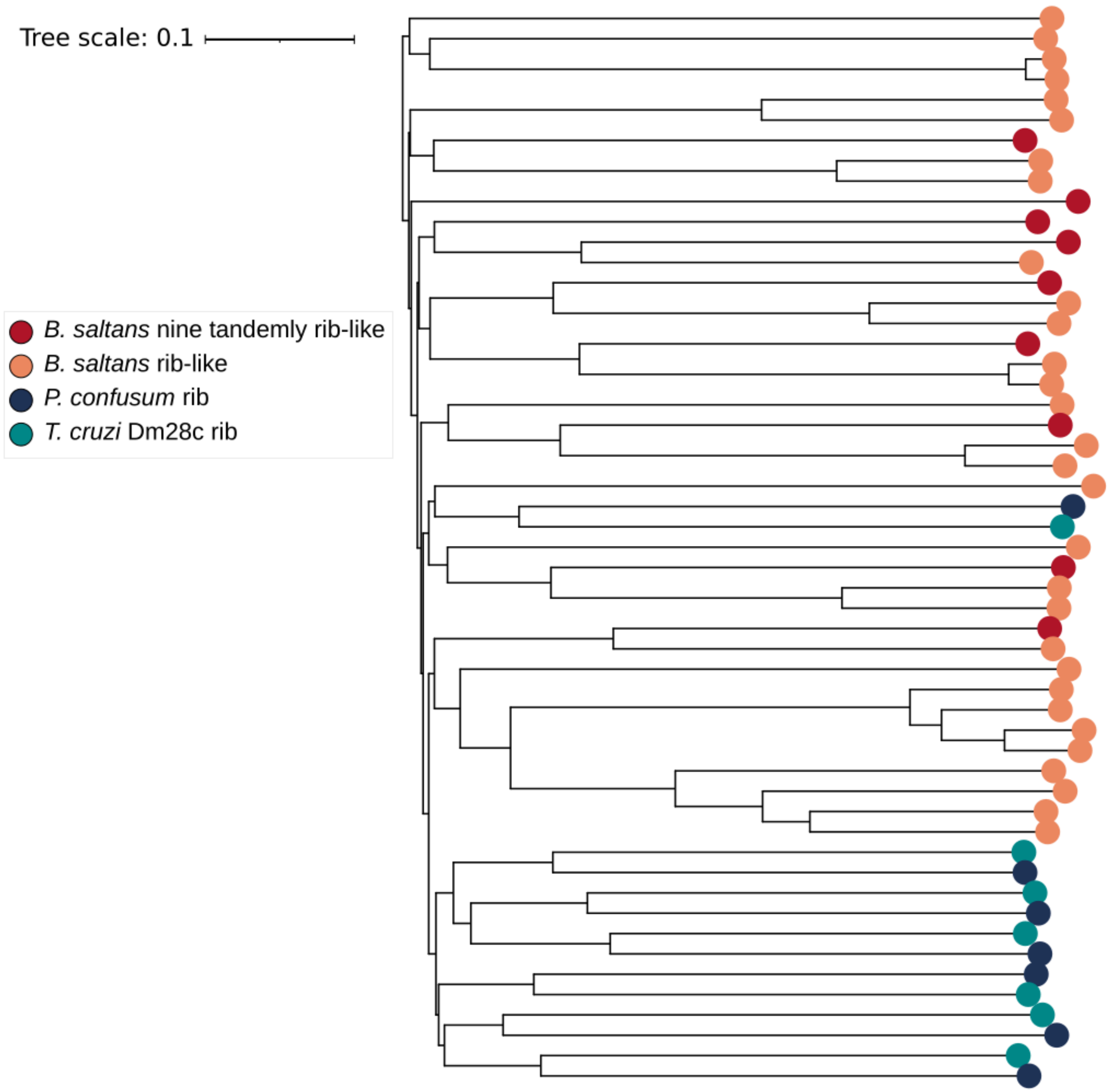


**Supplementary Figure S9. Structural relationships among DGF-1 ribs modules inferred with FoldTree.**

FoldTree was used to reconstruct the structural relationships among DGF-1 ribs modules of *Trypanosoma cruzi* Dm28c and *Paratrypanosoma confusum*, together with ribs-like regions identified in *Bodo saltans* proteins. Branch tips are color-coded according to species, as indicated in legend.

Figure S10

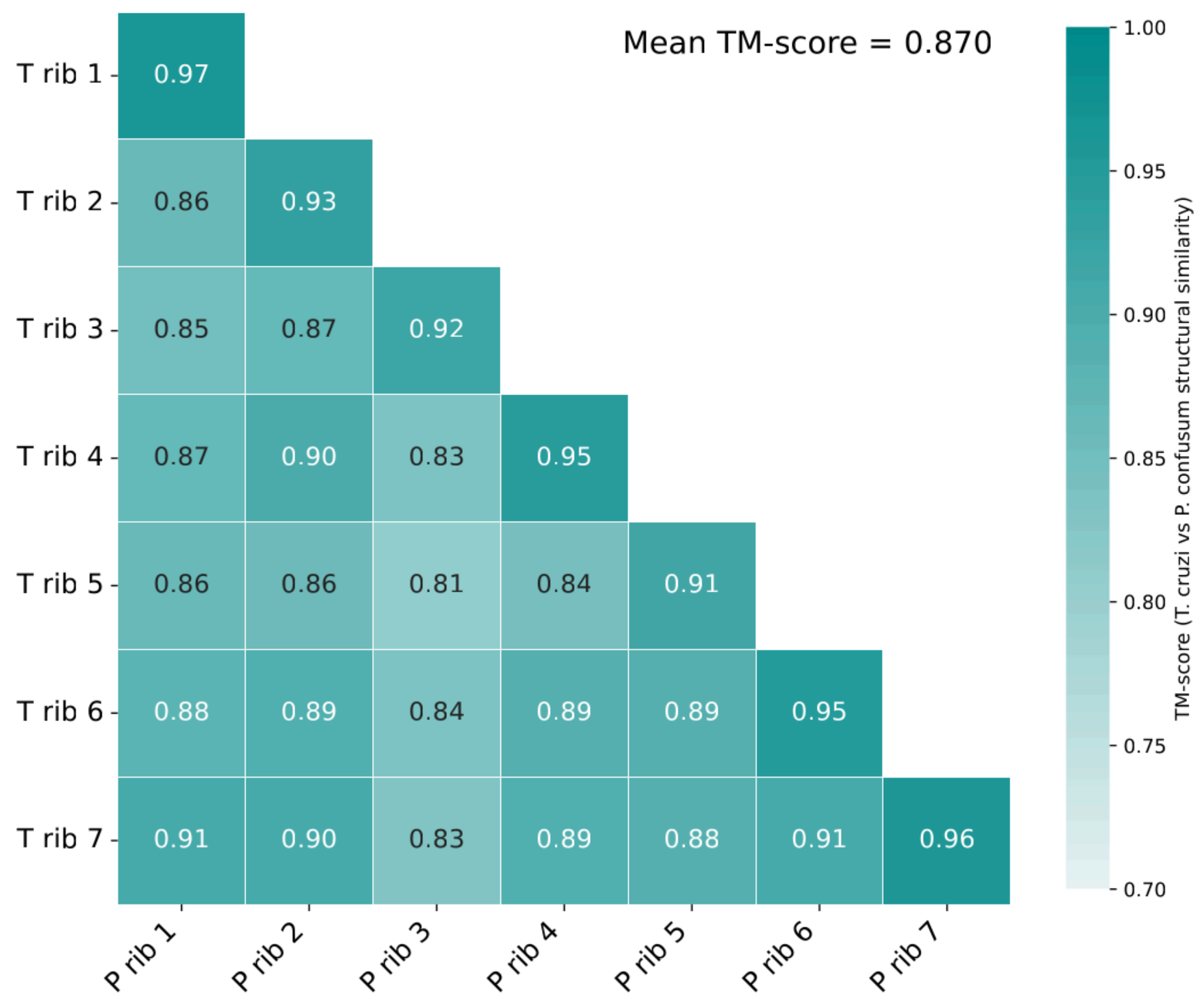


**Supplementary Figure S10**

Structural similarity matrix (Heatmap) based on cross-species pairwise TM-score comparisons between the individual rib modules of a representative full-length *T. cruzi* Dm28c architecture (y-axis) and those of *Paratrypanosoma confusum* (x-axis). Modules were modeled using AlphaFold3 and subjected to pairwise structural alignments.

Figure S11

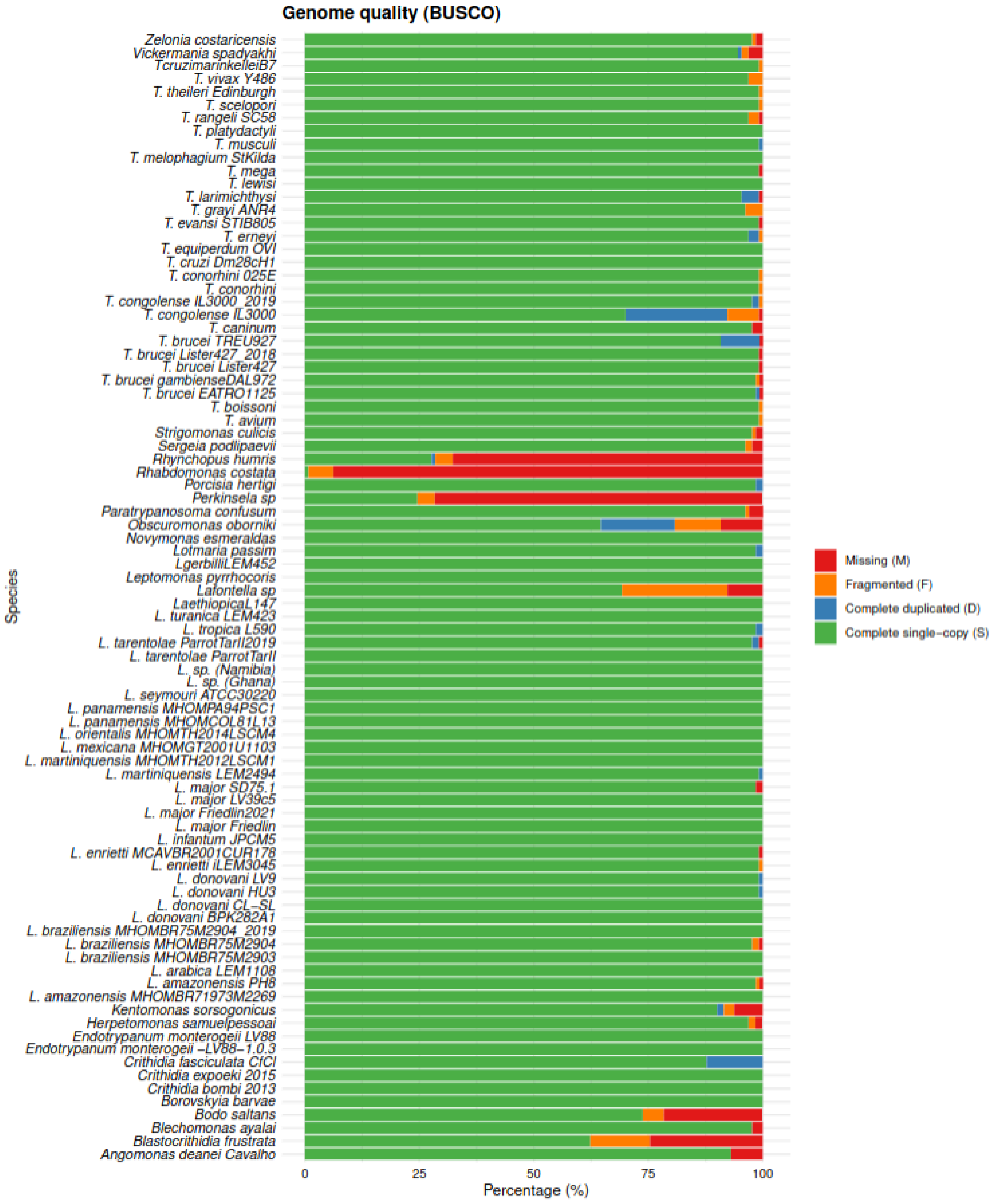


**Figure S11. BUSCO genome completeness estimated on the list of predicted genes.**

Genome completeness was evaluated using BUSCO v5.8.3 with the *euglenozoa_odb10* lineage dataset (n = 87 BUSCOs). Stacked bars show the proportions of complete single-copy (green), complete duplicated (blue), fragmented (yellow), and missing (red) BUSCO genes for each predicted proteome.

### Figure S12

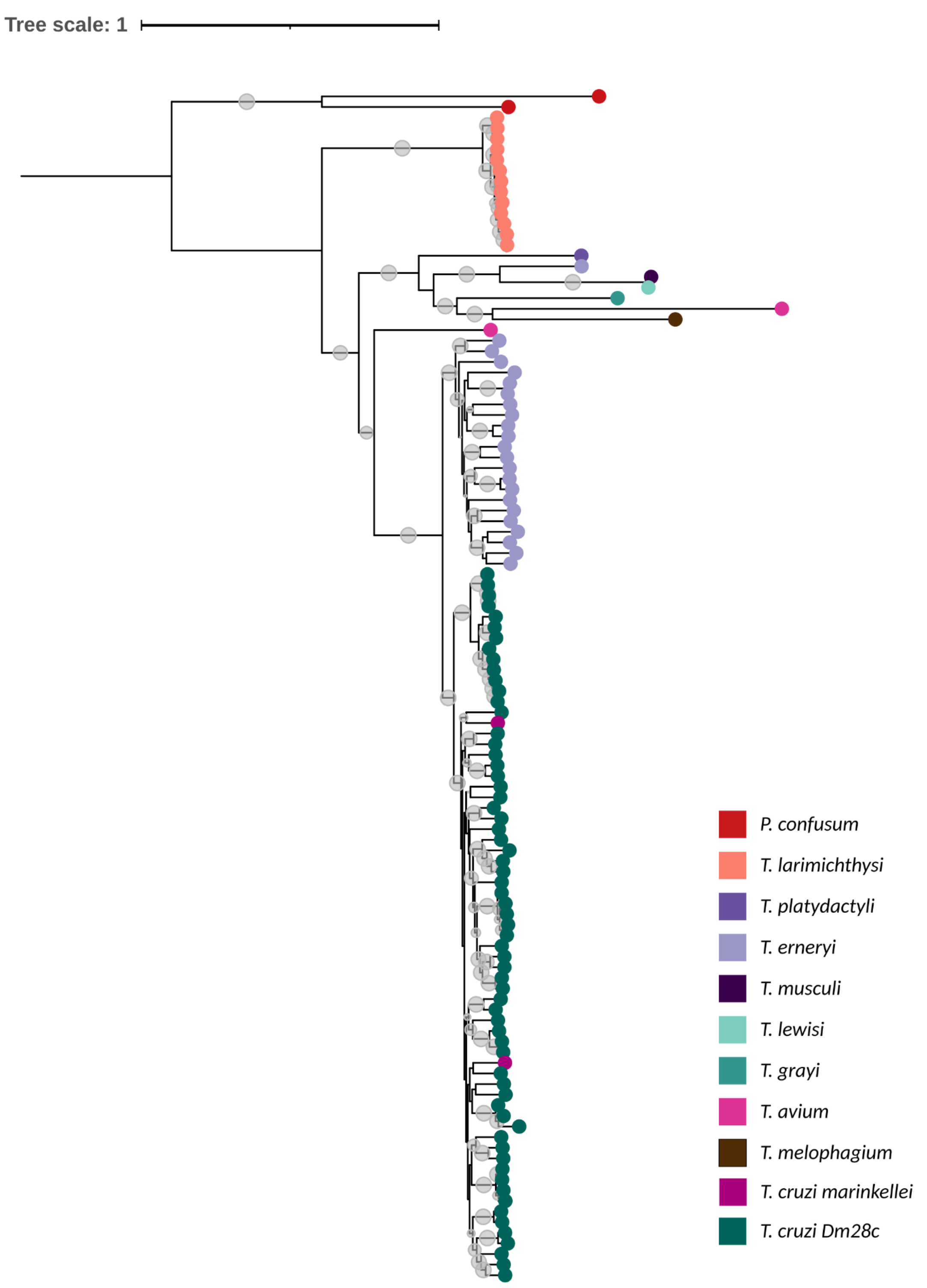
Tree scale: 1
P. confusum
T. larimichthysi
T. platydactyli
T. erneryi
T. musculi
T. lewisi
T. grayi
T. avium
T. melophagium
T. cruzi marinkellei
T. cruzi Dm28c

**Supplementary Figure S12. Phylogenetic relationships of DGF-1 homologues among Kinetoplastids.**

Rooted Maximum-likelihood phylogeny reconstructed from full-length DGF-1 amino acid sequences identified across DGF-1 containing species. The tree was rooted using the DGF-1 homolog from *Paratrypansoma confusum*. Branch tips are colored according to species, as indicated in the legend. The gray circle indicated branches with ultrafast bootstrap support values ≥ 75%.
